# Increased Cholesterol Synthesis Drives Neurotoxicity in Patient Stem Cell-Derived Model of Multiple Sclerosis

**DOI:** 10.1101/2024.01.16.575826

**Authors:** Rosana-Bristena Ionescu, Alexandra M. Nicaise, Julie A. Reisz, Eleanor C. Williams, Pranathi Prasad, Monika Dzieciatkowska, Daniel Stephenson, Marta Suarez Cubero, Liviu Pirvan, Cory M. Willis, Luca Peruzzotti-Jametti, Valentina Fossati, Frank Edenhofer, Tommaso Leonardi, Christian Frezza, Irina Mohorianu, Angelo D’Alessandro, Stefano Pluchino

**Affiliations:** Department of Clinical Neurosciences and NIHR Biomedical Research Centre, University of Cambridge, Cambridge CB2 0AH, UK; Department of Biochemistry and Molecular Genetics, University of Colorado School of Medicine, Aurora, CO, 80045 USA; Wellcome-MRC Cambridge Stem Cell Institute, Jeffrey Cheah Biomedical Centre, University of Cambridge, Cambridge CB2 0AW, UK; Genomics, Stem Cell Biology and Regenerative Medicine Group, Institute of Molecular Biology & CMBI, Leopold-Franzens-University Innsbruck, Innsbruck, Austria; Department of Metabolism, Digestion and Reproduction, Imperial College London, United Kingdom; The New York Stem Cell Foundation Research Institute, New York City, NY, 10019 USA; Center for Genomic Science of IIT@SEMM, Instituto Italiano di Tecnologia (IIT), 20139 Milan, Italy; University Hospital Cologne, Cologne, Germany

**Keywords:** Neuroimmunology, multiple sclerosis, neural stem cells, cellular senescence, cholesterol metabolism, lipid droplets, multi-omics, SASP, neurotoxicity, disease modelling

## Abstract

Senescent neural progenitor cells have been identified in brain lesions of people with progressive multiple sclerosis (PMS). However, their role in disease pathobiology and contribution to the lesion environment remains unclear.

By establishing directly induced neural stem/progenitor cell (iNSC) lines from PMS patient fibroblasts, we studied their senescent phenotype *in vitro*. Senescence was strongly associated with inflammatory signaling, hypermetabolism, and the senescence associated secretory phenotype (SASP). PMS-derived iNSCs displayed increased glucose-dependent fatty acid and cholesterol synthesis, which resulted in the accumulation of cholesteryl ester-enriched lipid droplets. An HMG-CoA reductase-mediated lipogenic state was found to induce secretion of the SASP in PMS iNSC conditioned media via transcriptional regulation by cholesterol-dependent transcription factors. SASP from PMS iNSCs induced neurotoxicity. Chemical targeting of HMG-CoA reductase using the cholesterol-lowering drug simvastatin (SV) prevented SASP release and resulting neurotoxicity.

Our findings suggest a disease-associated, cholesterol-related, hypermetabolic phenotype of PMS iNSCs that leads to neurotoxic signaling and is rescuable pharmacologically.

## INTRODUCTION

Multiple sclerosis (MS) affects 2.8 million individuals worldwide, which makes it the most common inflammatory autoimmune demyelinating disease of the central nervous system (CNS).^1^ Progressive MS (PMS) represents the late stage of the disease and is marked by a steady and unrelenting accumulation of chronic neurological disability as the clinical outcome of continuing neuronal loss.^2^ In PMS several immune-dependent and -independent mechanisms of disease have been described over the years.^3^ However, there are very little to no disease-modifying therapies (DMTs) for PMS, as the complexity of this disease has made it particularly challenging to identify targets which promote neuroprotection.^3^ Hence, the identification of key mechanisms of disease, and the development of a new class of DMTs capable of effectively promoting neuroprotection to slow disease progression, are critical unmet needs for people with PMS.^3^

PMS shares many pathological hallmarks with classic neurodegenerative diseases (including Alzheimer’s disease and Parkinson’s disease)^4^, with age being the primary risk factor for the progression of disease.^5^ Longitudinal analysis of the brain captured with magnetic resonance imaging (MRI) in people with MS revealed signatures indicative of advanced brain aging compared with healthy individuals and patients diagnosed with other neurodegenerative disorders, such as Parkinson’s disease and dementia.^6,7^

As individuals undergo biological aging, so do their cells. Cellular aging or senescence is characterized both by the gain-of-function of maladaptive processes and the loss-of-homeostatic processes, the combination of which disrupts normal cellular functions^8^. Multiple, independent post-mortem pathological studies of human individuals at advanced age and patients with neurodegenerative diseases have shown an accumulation of senescent cells in the brain.^9,10^ Senescent neural stem cells (NSCs) were also identified within lesions of PMS brains.^11^ Moreover, human single-nuclear sequencing analyses of the post-mortem MS brain^12^ and genome-wide association studies (GWAS) have identified transcriptional states and genetic variants indicative of intrinsic cell dysfunctions associated with altered cellular energetics and senescent states.^13,14^ The impact of dysfunctional cellular responses, such as senescence and altered energetics, as well as the potential role for accelerated brain age, neuronal function, and reduction of the neurocognitive reserve^7,15^ is yet to be established in PMS.

Reprogramming technologies, including reverting somatic cells to an embryonic-like state (induced pluripotent stem cells - iPSCs) or direct conversion to different cell types, are being extensively applied to establish disease-in-a-dish human models of CNS disorders that cannot be fully recapitulated by animal models. iPSC-derived CNS cells (such as neurons, oligodendrocytes, astrocytes, microglia, and endothelial cells) have been effectively used to model intrinsic cellular dysfunctions of MS. By studying these systems, abnormalities have been found in mitochondrial function, junction integrity, and the development of a (resting) inflammatory state.^16–20^ Senescent features have been described in iPSC-NSCs derived from people with PMS by identifying a SASP that was found to inhibit oligodendrocyte progenitor cell differentiation into mature myelinating cells.^11^

Cellular metabolism is a key component of senescence and disease-associated mechanisms. Hypermetabolism, or an abnormally high metabolic rate, has been described in erythrocytes^21^, fibroblasts^22^, and astrocytes^17^ from people with MS, as well as in the context of physiological aging.^23^ Analysis of cerebrospinal fluid (CSF) from individuals with MS revealed dysregulated metabolic features in glucose and energy pathways, amino acid metabolism, and lipid formation and breakdown.^24^ Lipid metabolism is involved in controlling neural stem/progenitor cell homeostasis and is linked as a major regulator of cellular senescence and the SASP.^25,26^ Whether an altered metabolic state is present in senescent NSCs from individuals with PMS is unknown.

Herein, we generated directly induced NSCs (iNSCs) from people with PMS (PMS iNSCs) and unaffected controls.^27^ We chose to use the direct induction of fibroblasts into NSCs as this method has improved fidelity in maintaining the aging signature of the donor.^28^ This affords the capability to more faithfully model disease- and age-associated metabolic alterations in PMS iNSCs and their functional implications. Using this model, we performed a multi-omics phenotypical characterization of cells, including intra and extracellular proteomics, metabolomics, and lipidomics. These data were then integrated with functional stable isotope tracing to uncover the key drivers of altered function in PMS iNSC.

We observed that PMS iNSCs exhibit a senescent gene expression signature and hypermetabolism *in vitro*. We identified lipid droplet (LD) accumulation as a key pathological hallmark and show that this is a cell-autonomous feature primarily emerges from glucose-derived, 3-hydroxy-3-methylglutaryl-CoA reductase (HMGCR) activity-dependent cholesterol synthesis. We next demonstrated that the increased HMGCR activity in PMS iNSCs regulates the paracrine neurotoxic SASP through transcriptional regulation by cholesterol-dependent transcription factors (TFs). Lastly, we provide evidence that the paracrine neurotoxic SASP of PMS iNSCs can be prevented by treatment with the HMGCR inhibitor simvastatin.

Our findings elucidate how a hypermetabolic state fueling cholesterol biosynthesis transcriptionally regulates the neurotoxic SASP in iNSCs from people with PMS. This highlights the potential contribution of NSC metabolism to the progression of disability and accelerated neurodegeneration observed in the PMS brain.

## RESULTS

### PMS iNSCs show senescent gene expression linked to inflammation and hypermetabolism

To model disease associated features of NSCs we applied cell reprogramming technologies. We first confirmed previous findings of senescence in PMS NSCs^11^ by generating iPSCs from skin-derived fibroblasts (**Fig. S1A-B, Table 1**). We derived endodermal, mesodermal, and ectodermal iPSC progenies (**Fig. S1C**), and found significantly increased expression of the cyclin-related cellular senescence marker *CDKN2A*, but not *CDKN1A* or *TP53*, in the PMS ectodermal progenies only (**Fig. S1D**). Hence, the acquisition of an ectodermal NSC identity in PMS cells is associated with increased expression of senescence-associated genes in a cell-autonomous fashion.

**Table 1.** Cells and Cell lines.

| **Code** | **Sex** | **Age**<br>*(at collection)* | **Ethnicity** | **Condition** | **Cell Lines Generated** |
| --- | --- | --- | --- | --- | --- |
| Control A | Male | 39 | N/A | HC | iNSCs, fibroblasts |
| Control B | Female | 25 | N/A | HC | iNSCs, fibroblasts |
| Control C | Male | 28 | Caucasian | HC | iNSCs; fibroblasts; iPSCs; iPSC-derived endo-, meso- and ectoderm |
| Control D (WIBJ) | Female | 55-59 | White British | HC | iPSCs; iPSC-derived endo-, meso- and ectoderm |
| PMS A | Male | 56 | Caucasian | PPMS | iNSCs; fibroblasts; iPSCs; iPSC-derived endo-, meso- and ectoderm |
| PMS B | Male | 42 | African American | SPMS | iNSCs; fibroblasts |
| PMS C | Male | 46 | Caucasian | SPMS | iNSCs; fibroblasts |
| PMS D | Female | 63 | Caucasian | SPMS | iNSCs; fibroblasts; iPSCs; iPSC-derived endo-, meso- and ectoderm |

N/A: not available; HC: Healthy control; iNSC: induced neural stem cell; iPSC: induced pluripotent stem cell; PPMS: Primary progressive multiple sclerosis; SPMS: Secondary progressive multiple sclerosis.

As iPSC reprogramming has been demonstrated to rejuvenate the epigenome^29^, we next utilized direct reprogramming technology to generate cell lines that retain aging-related markers.^28^ We reprogrammed skin fibroblasts from 3 non-diseased individuals (Ctrl) and 4 people with PMS between 25 to 63 years of age into iNSCs (**Table 1**).^27,30^ Quality control analysis by immunocytochemistry showed superimposable expression of the established NSC/radial glial markers NESTIN, SOX2, and ETNPPL in both Ctrl and PMS iNSC lines (**Fig. 1A**).^28^ RT-PCR of *NESTIN*, *SOX2*, and *PAX6* confirmed comparable expression levels via band density across all iNSC lines, while the pluripotency marker gene *OCT4* was absent (**Fig. S1E**). We confirmed full viral clearance by assessing the Sendai virus vectors *KOS* and *C-myc*, as well as the viral genome using *SeV* targeting primers (**Fig. S1E**).

**Figure 1.**
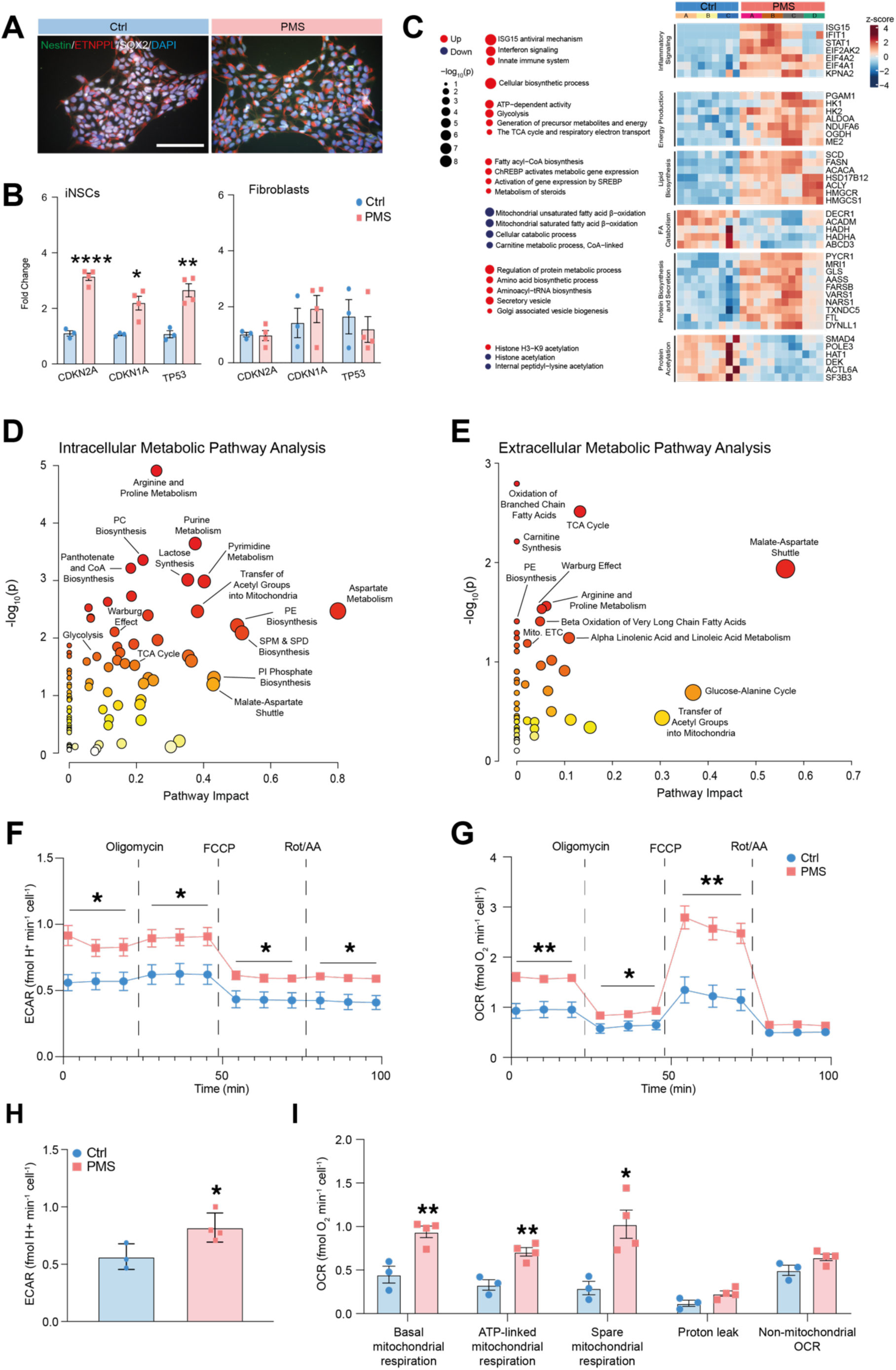
PMS iNSCs exhibit senescence associated inflammatory signaling and hypermetabolism. (**A**) Immunocytochemistry of accepted NSC markers in Ctrl and PMS iNSCs. Scale bar: 50 μm. (**B**) mRNA expression of senescence-associated genes. (**C**) Pathway enrichment analysis of GO and REACT terms in PMS *vs.* Ctrl iNSCs based on intracellular proteomics. Selected proteins from pathways are plotted in the accompanying heatmap. Differentially enriched proteins and pathways were considered significant at adjusted p-value< 0.05 (Benjamini-Hochberg correction) under edgeR and hypergeometric tests, respectively. (**D-E**) Metabolic Pathway Analysis of intracellular (**D**) and extracellular (**E**) metabolites. Differentially abundant metabolites were considered significant at adjusted p-value< 0.05 (Benjamini-Hochberg correction) under Welch’s t-test. (**F**-**G**) Extracellular acidification rate (ECAR, **F**) and oxygen consumption rate (OCR, **G**) over time during a mitochondrial stress test in the presence of glucose, pyruvate, and glutamine. (**H**) Basal ECAR quantified as in **F**. (**I**) Quantification of basal mitochondrial respiration, ATP-linked mitochondrial respiration, spare mitochondrial respiration, mitochondrial respiration linked to proton leak, and non-mitochondrial OCR using the mitochondrial stress test. Experiments in **B-I** were done on n= 3 Ctrl and n= 4 PMS iNSC lines, each performed in n ≥ 3 replicates. Data in **B**, **F**-**I** are as mean values ± SEM. *p≤ 0.05, **p≤ 0.01, unpaired t-test.

The expression of *CDKN2A*, *CDKN1A*, and *TP53* was significantly increased in PMS iNSC lines (*vs*. Ctrl iNSCs) (**Fig. 1B**).^28^ We observed no difference in senescence gene expression when comparing the parental PMS and Ctrl fibroblasts (**Fig. 1B**). These findings support the use of PMS iNSCs as a viable *in vitro* cell-modelling platform to explore pathobiology-relevant or -associated mechanisms of disease.

We next performed a mass spectrometry based untargeted intracellular proteomics analysis to uncover senescence-associated dysfunctional pathways. Principal component analysis (PCA) showed a separation between groups (**Fig. S1F**). Protein set enrichment analysis of up and downregulated proteins in PMS iNSCs (*vs.* Ctrl iNSCs) revealed biological process terms broadly falling within two categories: (1) inflammatory signaling and (2) metabolic processes related to energy production and anabolic/catabolic balance (**Fig 1C**). Upregulated proteins pertaining to inflammatory pathways were primarily related to interferon (IFN) signaling (ISG15, STAT1). Concordantly, western blot analysis confirmed significantly increased expression of ISG15, STAT1, and active form phosphorylated (p)-STAT1 (Y701) in PMS iNSCs (**Fig. S1G**). Upregulated proteins linked to cell metabolism in PMS iNSCs were found to be involved in energy production (glycolysis and mitochondrial respiration), lipid biosynthesis (sterols and fatty acids [FAs]), and protein biosynthesis and secretion (**Fig. 1C**). Downregulated proteins were associated with FA catabolism and protein acetylation (**Fig. 1C**). Overall, the proteomic analysis highlighted specific inflammatory and metabolic biological processes in PMS iNSCs that are linked to cellular senescence.^31–33^

As changes in protein expression are not necessarily predictive of differences in cellular metabolism, we next sought evidence of metabolic rewiring in the PMS iNSCs via mass spectrometry-based analysis of the intra- and extracellular metabolome. PCA confirmed a separation of both the intra- and extracellular metabolomes of Ctrl and PMS iNSC lines (**Fig S2A-B**). We identified several enriched metabolic pathways (**Fig. 1D-E**). In the intracellular metabolome, these included energy production through glycolysis and mitochondrial respiration (e.g., TCA cycle), lipid biosynthesis (e.g., transfer of acetyl groups into mitochondria), and amino acid metabolism (e.g., arginine and proline) (**Fig. 1D**). The energy production pathway was also reflected in the extracellular metabolome (e.g., TCA cycle and mitochondrial electron transport chain [ETC]) (**Fig. 1E**). These findings are in line with the metabolic landscape typical of senescent cells, including energetic hypermetabolism.^26^

We functionally validated our findings by assessing cellular bioenergetics using the Seahorse mitochondrial stress test. We found that PMS iNSCs have a significantly altered bioenergetic profile *vs.* Ctrl iNSCs (**Fig. 1F-G**) that is absent in the parental PMS and Ctrl fibroblasts (**Fig. S2C-E**). This altered bioenergetic profile was reflected in the increased basal extracellular acidification rate (ECAR), basal mitochondrial respiration, ATP-linked mitochondrial respiration, and spare mitochondrial respiratory capacity in PMS *vs.* Ctrl iNSCs (**Fig. 1H-I**). Additionally, the comparable OCR/ECAR ratio between the PMS and Ctrl iNSCs suggests that increased glycolysis is directly coupled to increased oxidative phosphorylation (OXPHOS) in PMS iNSCs (**Fig. S2F**).

Next, we profiled cellular mechanisms underpinning the bioenergetic abnormalities of PMS iNSCs. We first confirmed that the hypermetabolic state of PMS iNSCs was not the result of a differences in cell size (**Fig. S2G**). We next analyzed the mitochondrial phenotypes of the cells as potential drivers of senescence-associated hypermetabolism. We found no significant differences in mitochondrial content (**Fig. S2H-I**), mitochondrial copy number (**Fig. S2J**), and mitochondrial mass (**Fig. S2K**) in Ctrl and PMS iNSCs. Similarly, no overt differences in the mitochondrial network morphology (**Fig. S2H,L-N**), membrane potential (**Fig. S2O**), and superoxide production (**Fig. S2P**) were seen in Ctrl and PMS iNSCs.

These data suggest that intrinsic mitochondrial changes were not responsible for the observed hypermetabolic state of PMS iNSCs, which instead is linked with senescence-associated expression signatures related to inflammatory signaling and functional metabolic rewiring.

### Glucose fuels increased synthesis of lipid precursors in PMS iNSCs

To further investigate this hypermetabolic state, we next looked at the potential extrinsic drivers and focused on differences in mitochondrial substrate utilization *in vitro*. Ctrl and PMS iNSCs cultured for 24 hours in basal media actively consumed glucose and pyruvate, which was coupled with the production of glutamine (**Fig. 2A**). However, extracellular glucose levels were significantly decreased in PMS iNSCs, while glutamine and pyruvate levels were comparable between groups (**Fig. 2A**). Consistent with these findings, the product of glycolysis, extracellular lactate, was significantly higher in PMS iNSCs (**Fig. 2A**).

**Figure 2.**
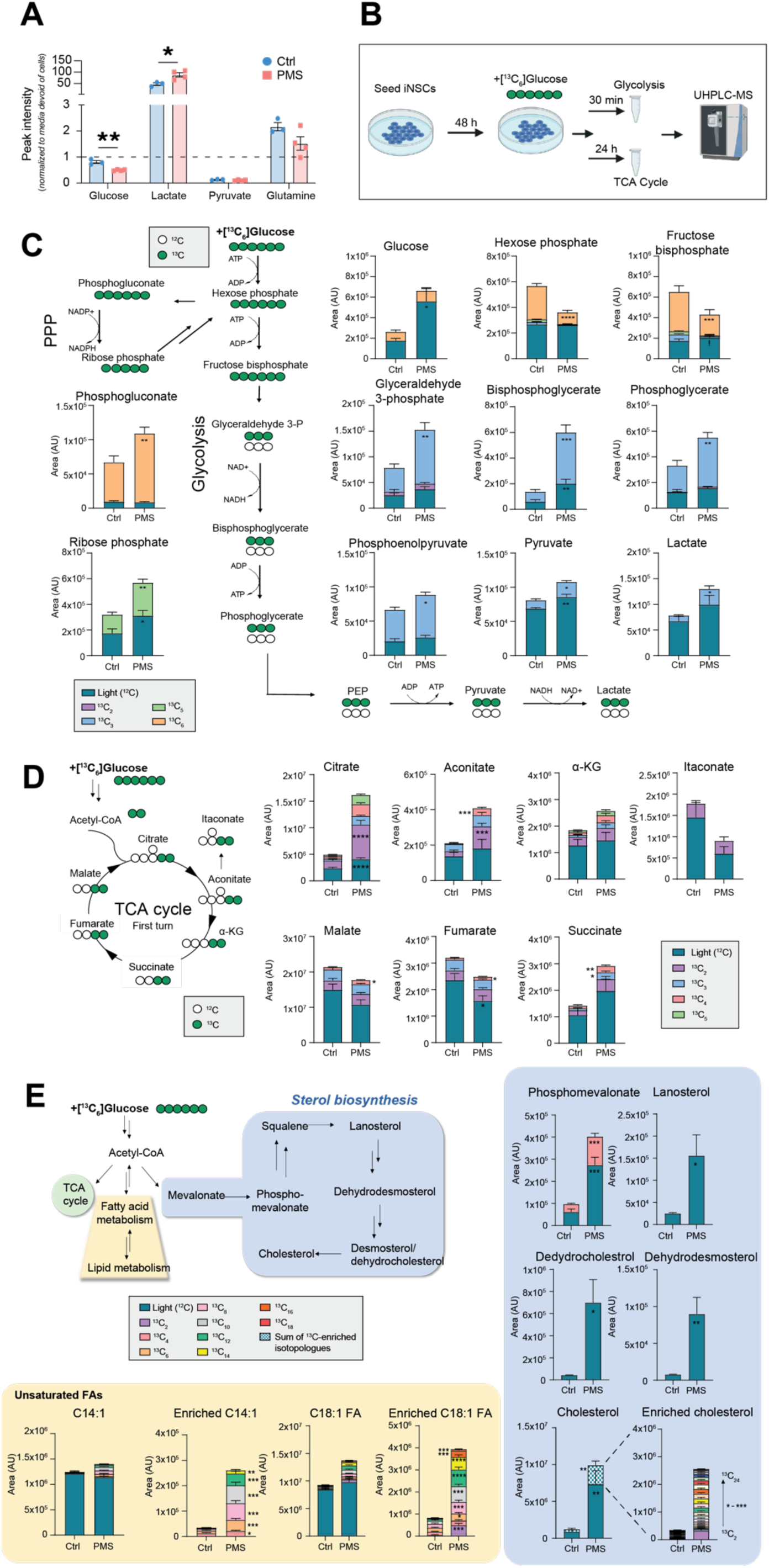
PMS iNSCs utilize glucose to support increased synthesis of lipid precursors. (**A**) Relative concentrations of glucose, lactate, pyruvate, and glutamine detected by UHPLC-MS in conditioned media. Values plotted over unconditioned media represented by dashed line. (**B**) Experimental schematic for [^13^C_6_]glucose tracing to assess glycolysis and tricarboxylic acid (TCA) cycle. (**C**) Intracellular enrichment in glycolysis at 30 min following [^13^C_6_]glucose tracing as in **B**. PPP, pentose phosphate pathway. (**D**) Intracellular enrichment in TCA cycle at 24 h following [^13^C_6_]glucose tracing as in **B**. (**E**) Intracellular enrichment in lipid biosynthesis pathways at 24 h following [^13^C_6_]glucose-tracing as in **B**. Experiments were done on n= 3 Ctrl and n= 4 PMS iNSC lines, each performed in n ≥ 3 replicates. Data in **A**, and **C-E** are mean values ± SEM. Stable isotope tracing data (**C**, **D** and **E**) is presented as peak areas (arbitrary units). *p≤ 0.05, **p≤ 0.01, ***p≤ 0.001, ****p≤ 0.001, unpaired t-test.

We then hypothesized that the observed metabolic phenotype may be the result of increased glucose utilization by PMS iNSCs. To test this, we employed serial stable isotope tracing experiments with [^13^C_6_]glucose to determine its intracellular fate (**Fig 2B**). At 30 min we found that glycolysis was more active in PMS iNSCs, as suggested by significant ^13^C_3_ enrichment of all mid and late glycolytic metabolites (from glyceraldehyde 3-phosphate through pyruvate and lactate) (**Fig. 2C**). Significantly increased levels of ^13^C_6_ 6-phosphogluconate and ^13^C_5_ ribose phosphate were also observed in PMS iNSCs, which supports higher glucose utilization through the pentose phosphate pathway (PPP) (**Fig. 2C**). At 24 hours, PMS iNSCs also showed an enrichment of ^13^C-labeled isotopologues of TCA intermediates (citrate, aconitate, α-ketoglutarate, succinate, fumarate, and malate), which is indicative of increased usage of glucose-derived carbons in the TCA cycle (**Fig. 2D**). These changes were associated with a higher percentage of labeled isotopologues as a fraction of total levels of TCA cycle metabolites **(Fig. S3A)**, which suggest an increased contribution of glucose-derived carbons to the TCA cycle in PMS iNSCs. However, the proportion of [^13^C_6_]glucose labeled TCA intermediates was found to decrease as the cycle progressed beyond α-ketoglutarate, implying an alternate carbon source for the lower segments of the TCA cycle (**Fig. S3A**).

To assess whether glutamine fueled the lower segment of the TCA cycle, we then performed [^13^C_5_]glutamine tracing also at 24 hours. Consistent with the labelled glucose tracing experiments, we observed elevated total levels of TCA cycle metabolites in PMS iNSCs (**Fig. S3B**). The proportions of [^13^C_5_]glutamine labeling in the TCA cycle were comparable between groups (**Fig. S3B**). As compared to the [^13^C_6_]glucose labeling, we found an increased proportion of [^13^C_5_]glutamine labeled TCA intermediates in the lower half of the TCA cycle (α-ketoglutarate-succinate-fumarate-malate) (**Fig. S3B**). These results suggest that in PMS iNSCs a fraction of the glucose-derived citrate exits the mitochondria into the cytosol whereas the glutamine-derived carbons preferentially supply the lower segments of the TCA cycle.

We next focused on understanding the fate of the increased pool of glucose-derived citrate that exited the mitochondria. We explored two possible pathways related to cytoplasmic citrate that was informed by the proteomics analysis described above: (i) protein acetylation and (ii) lipid biosynthesis.^34^ We compared global protein acetylation levels following [^13^C_6_]glucose labeling between groups and found few (4% of all mapped proteins) differentially acetylated (e.g., NOP14, HSPA8, ALYREF, POLR2E) (**Fig. S3C**). We concluded that protein acetylation is unlikely to be a major differential pathway for the glucose-derived citrate between the two groups. In contrast, [^13^C_6_]glucose tracing at 24 hours revealed a marked increase in the concentration of ^13^C labeled lipid precursors (**Fig. 2E, Fig. S3D**). We found that ^13^C enrichment in all FAs (from C10 through C18) was higher in PMS iNSCs, particularly in unsaturated FAs (representative plots shown for C14:1 and C18:1 FAs in **Fig. 2E**), compared to their saturated counterparts (representative plots for C:10:0, C12:0, C14:0, C18:0 in **Fig. S3D**). This suggests that the increased accumulation of unsaturated FAs in PMS iNSCs is caused by an increased de novo FA synthesis from glucose and FA desaturation.

Interrogation of the cholesterol biosynthesis pathway showed a marked activation in all PMS iNSC lines. This included a statistically significant increase in the total and ^13^C-enriched levels of the sterol precursor phosphomevalonate and significantly increased total levels of lanosterol, dehydrodesmosterol, and dehydrocholesterol in PMS *vs.* Ctrl iNSCs (**Fig. 2E**). Total cholesterol was also significantly elevated in PMS iNSCs along with levels of ^13^C enriched cholesterol. Consistent increases in cholesterol isotopologues in PMS iNSCs were observed across the range of ^13^C_2_ to ^13^C_24_ (**Fig. 2E**).

Thus, the metabolic rewiring of PMS iNSCs is linked with altered glucose and glutamine metabolism that ultimately leads to the increased biosynthesis of lipid precursors such as cholesterol and unsaturated FAs.

### Cholesterol synthesis leads to accumulation of lipid droplets and lipidome saturation

To investigate the fate of increased lipid precursor production in PMS iNSCs, we subjected iNSC pellets and their conditioned media (CM) to an untargeted mass spectrometry-based lipidomic analysis. We found that PMS iNSCs harbor a distinct intracellular lipid signature, with multiple lipid classes being increased (*vs.* Ctrl iNSCs; **Fig. 3A**). The lipid class that exhibited the biggest induction within the PMS iNSCs was cholesteryl esters (ChEs) with 70% lipid class involvement and a 12-fold change (**Fig. 3A**). A compositional analysis across intracellular lipid classes identified a higher proportion of shorter and more saturated acyl/alkyl groups in PMS iNSCs (**Fig. S4A**). Extracellularly, we found only minor differences between groups, with undetectable ChE levels (**Fig. S4B**, **Fig. 3A**). These data suggest that the increase in lipid precursors is accompanied by higher storage in PMS iNSC primarily in the form of ChEs. Accordingly, we found significantly increased lipid droplet (LD) accumulation in PMS iNSCs (**Fig. 3B**). Ultra-high-resolution imaging and 3D reconstruction revealed that most of the LD load of PMS iNSCs had a perinuclear localization (**Fig. S4C**). Overall, our results suggest that PMS iNSCs have a lipogenic state due to ChE accumulation associated with compositional changes in the lipidome.

**Figure 3.**
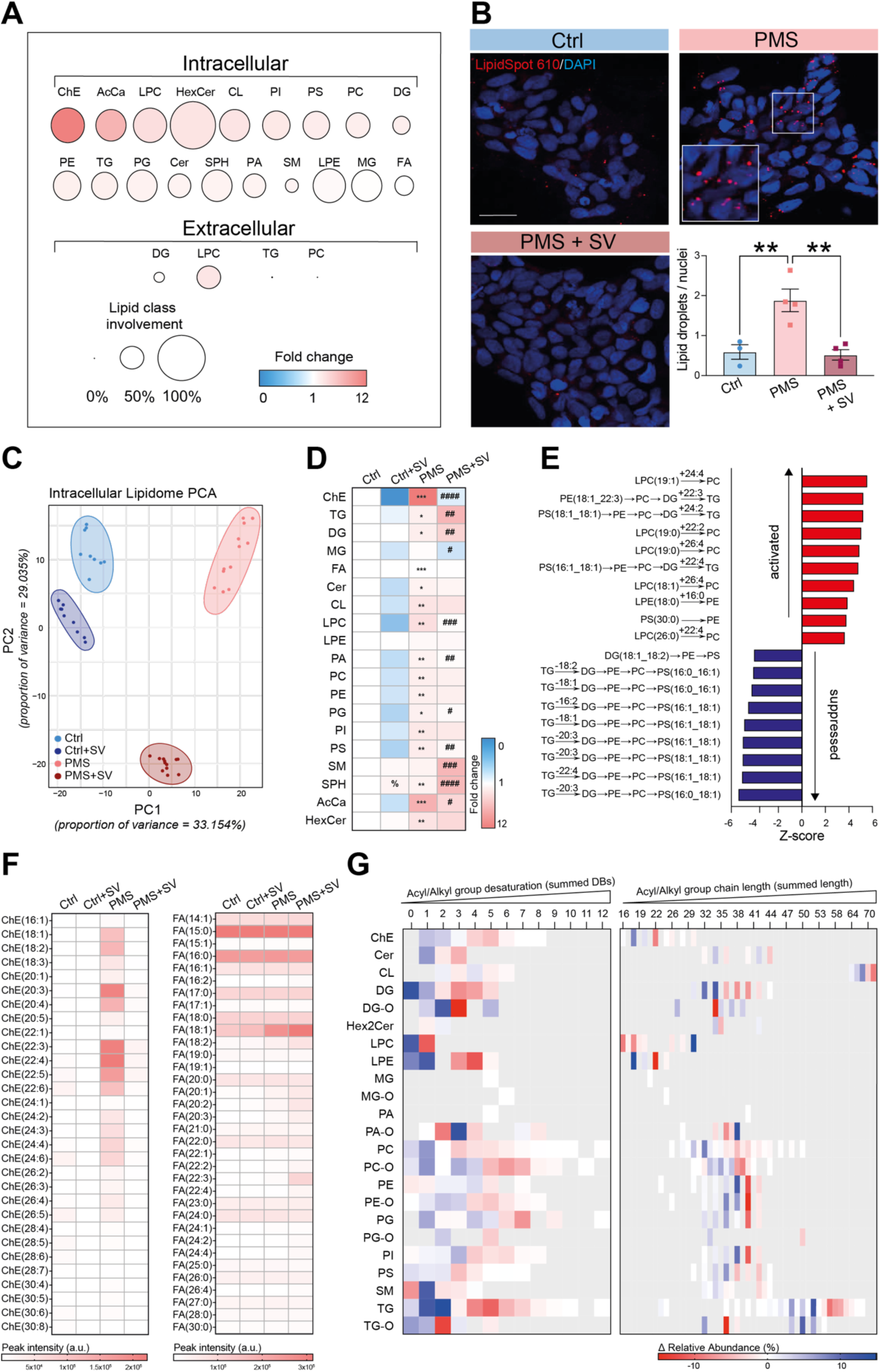
HMGCR pharmacological inhibition restructures the lipidome of PMS iNSCs. (**A**) Abundance and involvement of lipid classes in PMS *vs.* Ctrl iNSCs. Represented by fold change and proportion of differentially abundant lipid species within each lipid class. (**B**) Analysis of lipid droplet (LD) content. The bar graph shows epifluorescence-based quantification of LDs. Data are mean values ± SEM. Scale bar: 50 μm. SV= Simvastatin. **p≤ 0.01, one-way ANOVA, Tukey’s multiple comparisons. (**C**) PCA of the intracellular lipidome. (**D**) Fold changes in intracellular concentrations of lipid classes as in **C**. *p≤ 0.05, **p≤ 0.01, ***p≤ 0.001 in PMS *vs.* Ctrl; ^#^p≤ 0.05, ^##^p≤ 0.01, ^###^p≤ 0.001, ^####^p≤ 0.0001 in PMS *vs.* PMS+SV; ^%^p≤ 0.05 in Ctrl *vs.* Ctrl+SV; unpaired t-test. (**E**) BioPAN analysis of the top activated and suppressed intracellular lipid species reactions induced by SV treatment in PMS iNSCs. (**F**) Changes in intracellular concentrations of ChE and FA species as in **C**. Presented as peak areas (arbitrary units). (**G**) Structural changes in lipid species as in **E**. Relative abundance is expressed as percent change in acyl/alkyl group desaturation (as summed double bonds [DBs]) and in acyl/alkyl group chain length (as summed length). Experiments were done on n= 3 Ctrl and n= 4 PMS iNSC lines, each performed in n ≥ 3 replicates.

Given the accumulation of ChEs (**Fig. 3A**) and increased expression of the rate-limiting enzyme in cholesterol synthesis, 3-Hydroxy-3-Methylglutaryl-CoA Reductase (HMGCR) (**Fig. 1C**), we set out to test whether this pathway can be targeted to reduce the lipid load in PMS iNSCs. We chose to use the Food and Drug Administration (FDA)-approved oral medication simvastatin (SV) to pharmacologically inhibit HMGCR activity. Treatment of PMS iNSCs for 48 hours with SV (PMS+SV) was sufficient to reduce the LD content of PMS+SV iNSCs to Ctrl levels (**Fig. 3B, Fig. S4C**). Reducing LD accumulation in PMS+SV iNSCs induced a unique signature in the intracellular lipidome (**Fig. 3C**). Conversely, SV treatment of Ctrl iNSCs (Ctrl+SV) induced minor lipidome changes (**Fig. 3C**). ChEs were reduced to baseline Ctrl levels in PMS+SV iNSCs (**Fig. 3D**). Interestingly, PMS+SV had increased triglyceride (TG) levels compared to baseline PMS iNSCs (**Fig. 3D**). Given that LDs are primarily composed of TGs and ChEs, these results suggest that the accumulation of TGs is indispensable, while ChEs are responsible for inducing LD accumulation in PMS iNSCs.

Taken together these data show that PMS iNSCs are strong responders to SV treatment, which blocks the intracellular LD accumulation fueled by cholesterol biosynthesis and esterification.

We then hypothesized whether the compositional differences identified in the lipidome of PMS iNSCs, which extended beyond changes in ChEs, can be further explained by the disruption of pre-existing lipid class reactions involving LDs and the intracellular lipidome. Hierarchical clustering applied on all detectable lipid species suggested that SV treatment of PMS iNSCs resulted in the emergence of subsets of lipid species following four types of responses (**Fig. S4D**). Type 1 includes lipids that are different between groups (Ctrl *vs.* PMS) but are not affected by SV treatment. Type 2 includes lipids whose levels in PMS+SV iNSCs were restored to values observed in Ctrl iNSCs. Type 3 includes lipids that were found to be increased in PMS+SV iNSCs only (vs. all groups); while Type 4 includes lipids that were not affected by either the disease state (Ctrl *vs.* PMS) or treatment group (with or without SV). Notably, several lipid classes had species representation across all four response types (e.g., TG, PS [phosphatidyl serine], PC [phosphatidyl choline], DG [diglyceride], PE [phosphatidyl ethanolamine]). To understand why lipid species behaved differently in response to SV treatment we performed a deconvolution of lipid substructure using Linex^35^ software followed by hierarchical clustering. This analysis revealed a propensity of lipid species with higher summed acyl-chain lengths and lower saturation to follow a type 1 or type 3 response (**Fig. S4E**). Conversely, lipid species with lower summed acyl-chain lengths tended to follow a type 2 or type 4 response (**Fig. S4E**). These results suggest that the structural composition of lipids influences their response to HMGCR inhibition.

To further elucidate the mechanism underpinning the differential response of lipid species to SV treatment depending on their structural composition we performed network analysis at lipid class and species level in PMS and PMS+SV iNSCs using BioPAN^36^. We observed an activation of acylation reactions and a concomitant suppression of diacylation reactions involving non-ChE lipid classes (**Fig. S4F**), which is consistent with a redistribution of acyl-groups from ChEs in LDs to other lipid classes. Specifically, the utilization of acyl groups 24:4, 22:3, 24:2, 22:2, 26:4, 16:0 and 22:4 for acylation reactions was increased, while the loss of acyl groups 18:2, 18:1, 16:2, 18:1, 20:3, 20:3, 22:4, 20:3 in deacylation reactions was suppressed in PMS + SV iNSCs (**Fig. 3E**). We found that the identical acyl groups were highly engaged in the process of cholesterol esterification in PMS iNSCs, whereas only slight increases were observed in the related FA species (**Fig. 3F**). This suggests the presence of a dichotomic utilization of these FA species for either cholesterol esterification or acylation reactions by other lipid species in PMS iNSCs. Accordingly, HMGCR inhibition with SV adjusted the lipid compositional profile of PMS iNSCs by increasing the proportion of longer and less saturated acyl/alkyl groups (**Fig. 3G**).

Hence, the diversion of the PMS iNSC metabolome towards increased LD formation by ChE synthesis leads to an altered structural and compositional landscape of the intracellular lipidome towards increased saturation and shortened acyl chain moieties.

### Cholesterol-regulated TFs induce SASP in PMS iNSCs

Lipid-laden senescent cells have been reported to exhibit SASP^26^. In fact, we identified proteins associated with secretory pathways to be increased in the PMS iNSCs (**Fig. 1C**). To further investigate the functional implications of the ‘lipid laden’ phenotype of PMS iNSCs, we analyzed the secreted proteins in CM via untargeted mass spectrometry-based proteomics. SASP factors^37,38^, and/or MS-associated molecules (e.g. TIMP1, TIMP2, MMP2, EPHA4)^39,40^ were increased in in the CM of PMS iNSCs, (**Fig. 4A**).

**Figure 4.**
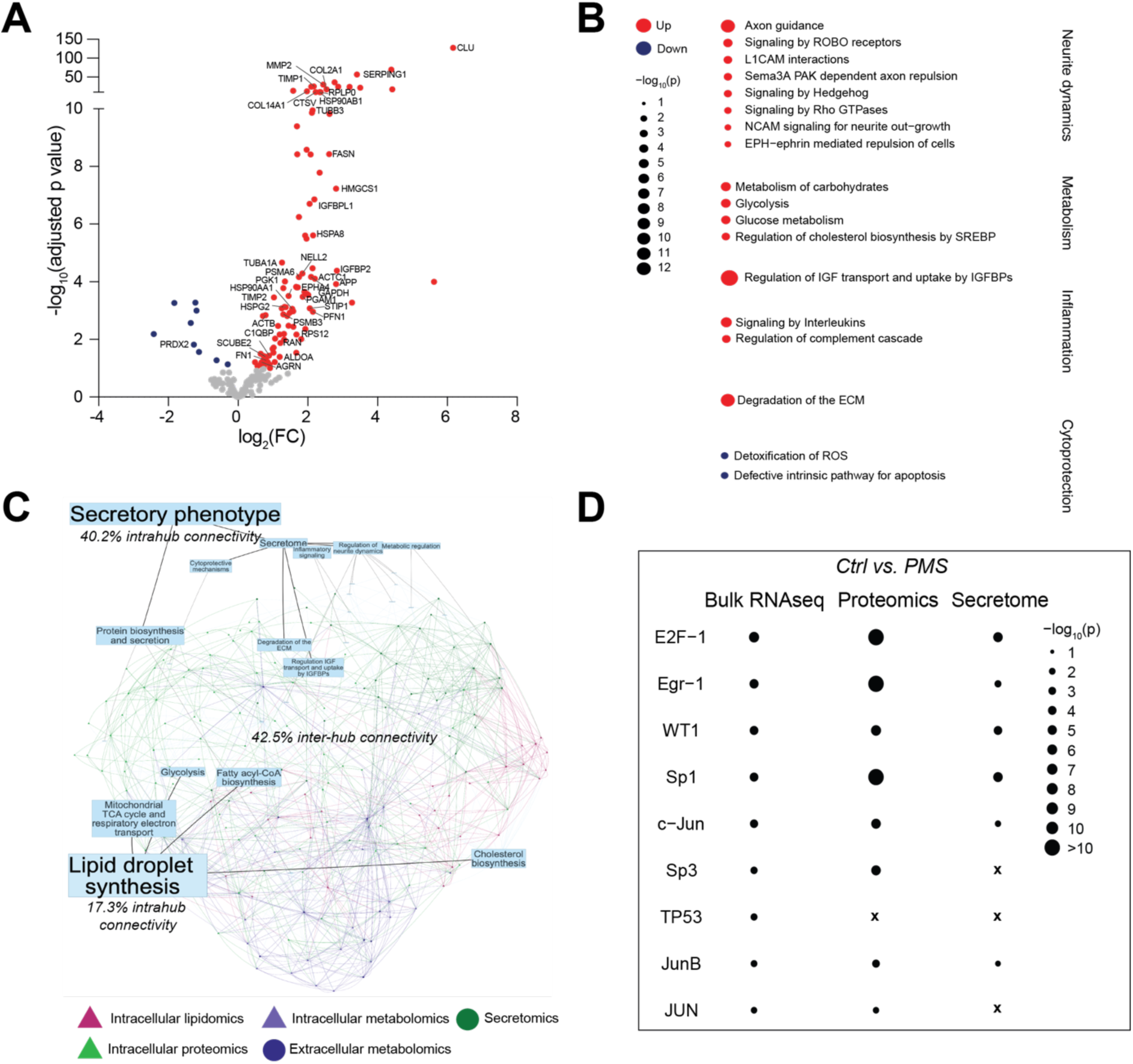
PMS iNSCs express cholesterol-regulated TFs that induce SASP. (**A**) Volcano plot of differentially secreted proteins (DSPs) in CM from PMS and Ctrl iNSCs. log_2_(FC) and adjusted p values are reported for each protein. (**B**) Pathway enrichment analysis of biological processes (GO and REACT terms) based on DSPs in CM from PMS and Ctrl iNSCs. Differentially enriched biological processes were considered significant at adjusted p≤ 0.05 under the hypergeometric test. (**C**) Covariation network Ctrl *vs*. PMS. Nodes are color-coded: proteins are represented in green, lipids in pink, metabolites in purple, with triangular nodes for intracellular and circles for extracellular metabolites or proteins. The strength of the edge between the nodes (proteins, metabolites, or lipids) represents the strength in the covariation. (**D**) Transcription factor (TF) enrichment analysis of TF terms abased on differentially expressed genes (DEGs) by BulkRNA Seq, proteins (DEPs) by intracellular proteomics analysis and DSPs as in **A**. DEGs were considered significant at adjusted p-value< 0.05 (Benjamini-Hochberg correction) under the DESeq2 test. DEPs were considered significant at adjusted p-value< 0.05 (Benjamini-Hochberg correction) under the edgeR test. TF term enrichment was considered significant at adjusted p≤ 0.05 under the hypergeometric test. X is for TF terms not being differentially enriched. Secretomics experiments were done on n= 3 Ctrl and n= 4 PMS iNSC lines, each performed in n ≥ 3 replicates. BulkSeq data analyzed was performed on n= 2 Ctrl and n= 2 PMS iNSC lines, each performed in n ≥ 2 replicates. DSPs in **A**-**D** were considered significant at adjusted p-value<0.1 (Benjamini Hochberg correction) and |log_2_FC|>0.2 under the edgeR test.

To infer the potential functional implications of the observed secretory phenotype, we performed a protein set enrichment analysis of the up and down-regulated differentially secreted proteins (DSPs) found in the CM. Enrichment analysis of PMS upregulated DSPs identified biological processes associated with five categories: (1) neurite dynamics, (2) metabolic processes, (3) regulation of insulin growth factor (IGF) signaling, (4) inflammatory signaling, and (5) degradation of the extracellular matrix (ECM) (**Fig. 4B**). Enrichment analysis of PMS downregulated DSPs, instead, pertained to cytoprotective mechanisms such as reactive oxygen species (ROS) detoxification and modulation of apoptosis (**Fig 4B**).

To establish a functional correlative line of evidence on a regulatory mechanism between the lipogenic state of PMS iNSCs and their secretory phenotype, we then performed an integrative analysis of the multiple-omics datasets. This included intra- and extracellular metabolomics, intracellular proteomics, secretomics, and lipidomics. The multi-omics dataset was explored using bulkAnalyseR^41^ interactive apps, which facilitated the integration of multiple datasets through combined multi-omics covariation networks contrasting samples from Ctrl and PMS iNSCs. Based on this analysis, we focused on two main biological processes, denoted as parental hubs, which were overrepresented across several of the omics datasets: lipid droplet biosynthesis and the secretory phenotype (**Fig. 4C**).

Underlying associated secondary biological processes were identified for each parental hub (e.g., cholesterol biosynthesis is a secondary subordinate hub to lipid droplet synthesis) (**Fig. 4C**). Lastly, altered features arising from overrepresentation analyses of differentially abundant/expressed metabolites/proteins corresponding to different profiled datasets were selected for inclusion as nodes of the covariation network (e.g., the differentially abundant metabolite cholesterol and the differentially expressed intracellular protein HMGCR were selected as nodes of the cholesterol biosynthesis secondary hub).

Focusing on the top 1000 connections (edges) between proteins, metabolites, and lipids, and further restricting the analysis to only entities linked to one of the central hubs, we found 57.5% pairs of connected entities were entirely contained within one hub (**Fig. 4C**). This included 40.2% (335 connections) within the secretory hub and 17.3% (153 connections) in the LD synthesis hub, compared to 42.5% (375 connections) of connections being formed between entities in opposite hubs (**Fig. 4C**). This revealed a dense connectivity between the LD synthesis hub and the secretory hub in PMS iNSCs (**Fig. 4C**). This high inter-hub connectivity by virtue of covariance may underly a functionally relevant regulatory link between the lipogenic state of PMS iNSCs and their functional paracrine signaling.

As the SASP is transcriptionally controlled^38,42,43^, we then performed a transcription factor (TF) enrichment analysis on differentially expressed mRNA species^28^, intracellular proteins, and secreted factors. We identified 6 significantly enriched TF candidates (E2F1, Egr1, WT1, Sp1, c-JUN, and JunB), which could account for the expression signatures observed in PMS iNSCs, across all three profiled expression modalities (**Fig. 4D**). Together, these TFs have strongly associated binding sites, trans modulatory functions, and cooperatively engage in gene regulatory networks^44–46^. These results point towards transcriptional co-modulation as a likely regulatory mechanism for the observed SASP of PMS iNSCs.

Evidence exists that the activity of several of the top identified TFs - which could account for the observed PMS iNSC secretome - is regulated by cholesterol (WT1^47^, c-JUN^48^, JunB^48^, Sp1^49^), or by the metabolic pathways of cholesterol synthesis (E2F1^50^), cholesterol esterification (E2F1^51^) or FA saturation (E2F1^52^). Hence, we propose that the lipogenic state and the secretory phenotype of PMS iNSCs are transcriptionally co-regulated.

### Pharmacological inhibition of HMGCR rescues the neurotoxic secretome of PMS iNSCs

To determine if cholesterol is regulating the SASP we subjected the CM of Ctrl+SV and PMS+SV iNSCs to untargeted mass spectrometry-based proteomics. We found more DSPs when comparing the CM of PMS *vs*. PMS+SV, compared to the CM of Ctrl *vs*. Ctrl + SV iNSCs (**Fig. 5A**). PMS+SV rescued the levels of previously identified DSPs in the PMS iNSC secretome (e.g., EPHA4, TIMP2, PRDX2) (**Fig. 5B**). Similar to the response of PMS iNSCs to SV treatment observed in the lipidome, we found an induction of novel DSPs in the CM of PMS+SV iNSCs. Some of the induced DSPs were unique and some were shared with the CM of Ctrl+SV. Several of these altered proteins were enzymes involved in glucose metabolism (e.g., PGK1, GAPDH, MDH1) and in cholesterol biosynthesis and esterification (e.g., HMGCS1, ACAT2, FASN), indicating a common compensatory response to HMGCR inhibition (**Fig. 5A-B**). Nevertheless, these DSPs reached much higher abundance in the CM of PMS+SV *vs*. the CM of Ctrl+SV iNSCs, suggesting an underlying regulatory mechanism as opposed to a pleiotropic effect of SV treatment as the cause for the observed secretory induction. This overall suggests that PMS iNSCs are strong responders to HMGCR inhibition with regards to their secretome.

**Figure 5.**
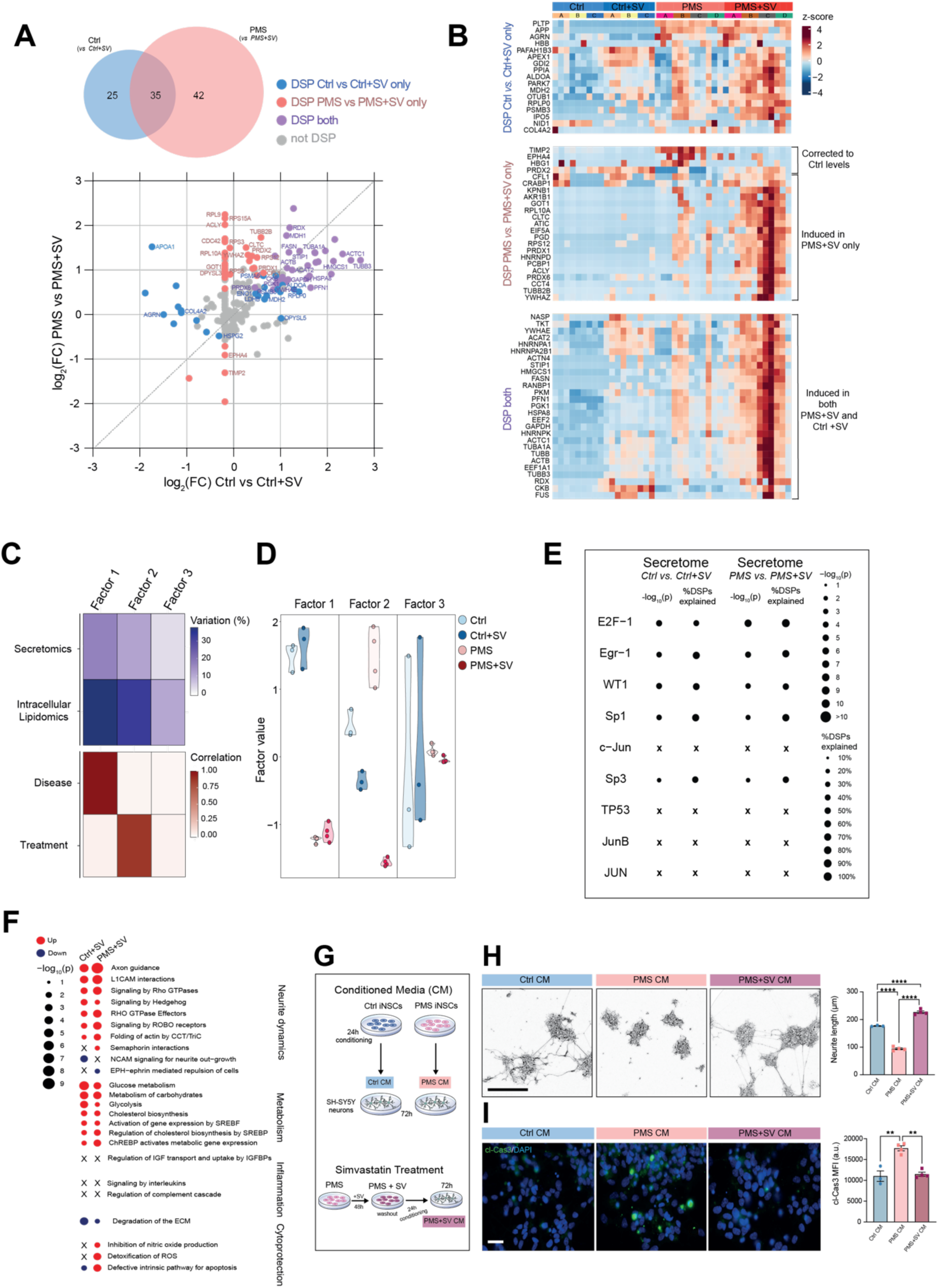
HMGCR pharmacological inhibition rescues the neurotoxic SASP of PMS iNSCs. (**A**) Venn diagram and cross plot of DSPs in non-treated and SV-treated groups of both Ctrl and PMS iNSCs. Log_2_(FC) are reported for each comparison. (**B**) Heatmap of DSPs based on relative z-scores, as in **A**. (**C**) Multi-Omics Factor Analysis (MOFA) of secretomics and intracellular lipidomics and correlation for calculated factors separated by disease or treatment. (**D**) Distribution of factor values within groups and by factors. (**E**) Transcription factor (TF) enrichment analysis based on identified DSPs, and proportion of total DSPs explained by each TF term, as in **A**. TF term enrichment was considered significant at adjusted p≤ 0.05 under the hypergeometric test. X is for TF terms not being differentially enriched. (**F**) Pathway enrichment analysis of biological processes (GO and REACT terms) based on SV-induced DSPs in CM from PMS and Ctrl iNSCs. Differentially enriched biological processes were considered significant at adjusted p≤ 0.05 under the hypergeometric test. X is for pathways not being differentially enriched. (**G**) Experimental schematic of CM and CM SV Treatment paradigms. (**H**) Representative phase contrast microscopy images and quantification of neurite length as in **G**. Scale bar= 250 μm. (**G**) Representative epifluorescence microscopy images and quantification of cleaved caspase-3 (cl-Cas3) expression as in **G**. Scale bar: 50 μm. Experiments were done on n= 3 Ctrl and n= 4 PMS iNSC lines, each performed in n ≥ 3 replicates. DSPs in **A**-**F** were considered significant at adjusted p-value<0.1 (Benjamini Hochberg correction) and |log_2_FC|>0.2 under the edgeR test. Differentially abundant lipids in **C** and **D** were considered significant at adjusted p-value< 0.05 (Benjamini-Hochberg correction) under Welch’s t-test. Data in **H** and **I** are mean values ± SEM. **p≤ 0.01, ****p≤ 0.0001, one-way ANOVA with Tukey’s multiple comparisons.

To integrate and summarize the observed variation between experimental groups, across modalities (intracellular lipidomics and secretomics), and propose biologically relevant regulatory interactions, we applied multi-omics factor analysis (MOFA)^53^. The MOFA model was used to fit latent factors on the intracellular lipidome (n= 642) and secretome (n= 241) from samples of untreated and SV-treated Ctrl and PMS iNSCs. The total percentage of variance explained by the factors was higher in the lipidome dataset than in the secretome dataset across all factors which is likely due in part to the larger number of features in this modality (**Fig. 5C**). Factors 1 and 2 explained a non-negligible percentage of variance in both-omic views, and thus could be seen as factors that are common to both lipidomic and secretomic modalities. Together, they explain 70.8% and 27.7% of the total variance in the lipidome and secretome modalities, respectively. To link factors 1 and 2 to disease state and treatment, we then assessed the relationship among the factors and the experimental groups (**Fig. 5C-D**).

Factor 1 was found to be strongly correlated with the disease state (Ctrl or PMS), which we identified as major driver of variation, and did not discriminate between untreated and SV-treated cells (**Fig. 5C-D**). Factor 1 loadings in the lipidome modality showed high weights for lipid species previously identified as nonreactive to SV treatment, as well as an enrichment in saturated acyl groups which are not involved in cholesterol esterification (**Fig. S5A**). Factor 1 loadings in the secretome modality showed high weights for DSPs between Ctrl and PMS iNSCs which were unaffected by SV treatment (e.g., IGFBP2, SERPING1, APP) (**Fig. S5B**), suggesting that these secretory features are unlikely to be regulated by the HMGCR-mediated lipogenic state of PMS iNSCs.

Instead, factor 2 was strongly associated with the treatment state (with or without SV) (**Fig. 5C-D**). This also captured the strong response to SV by PMS iNSCs compared to Ctrl iNSCs. Factor 2 loadings in the lipidome modality showed high weights for lipid species whose reactivity we previously identified as modified in response to SV treatment (**Fig. 3E and Fig. S5C**). We also noted an enrichment in unsaturated FA species involved in the increased cholesterol esterification in PMS iNSCs, as previously identified in the lipidomics analysis (**Fig. 3F and Fig. S5C**). Factor 2 loadings for the secretome modality showed high weights for DSPs between PMS and control iNSCs corrected by SV treatment (e.g., EPHA4, PRDX2), and DSPs de novo induced by SV treatment in PMS iNSCs (e.g., HMGCS1, ACAT2, FASN) (**Fig. S5D**), suggesting that these secretory features are likely regulated by the HMGCR-mediated hyper-lipogenic state of PMS iNSCs.

We next performed TF enrichment analysis using the DSPs from Ctrl *vs*. Ctrl + SV iNSCs and from PMS *vs*. PMS+SV iNSCs. We identified E2F1, Egr1, WT1, Sp1 and Sp3 as top enriched candidates, which account for most of the secretory signatures mediated by SV treatment in Ctrl and PMS iNSCs (40-60% and 50-90%, respectively) (**Fig. 5E**). Taken together, our results suggest that the HMGCR-mediated hyper-lipogenic state of PMS iNSCs regulates their secretory profile via transcriptional regulation by cholesterol-dependent TFs.

As statins have been reported to have either anti-senescence or senescence-inducing effects depending on the cellular context in which they act^54^, we also measured the effect of SV treatment on the expression of the senescence-associated genes CDKN2A, CDKN1A and TP53 as in **Fig. 1B**. Interestingly, SV had no effect on PMS iNSC expression of CDKN1A and TP53 transcripts, while significantly increasing the expression of CDKN2A (1.5-fold increase) (**Fig. S5F**). Hence, we concluded that the effects of SV/HMGCR inhibition on the PMS iNSC-associated SASP are independent of a direct modulatory effect on the senescence state itself.

We next explored which of the functional secretory signatures were affected by HMGCR inhibition. For this, we subjected previously identified DSPs in Ctrl iNSC CM *vs*. Ctrl+SV iNSC CM and PMS iNSC CM *vs*. PMS+SV iNSC CM to protein set enrichment analysis. HMGCR inhibition via SV treatment was able to correct some of the previously identified dysregulations in the PMS iNSC secretome related to the functional roles of EPH-Ephrin signaling, degradation of the ECM, and cytoprotective mechanisms such as ROS detoxification and modulation of apoptosis (**Fig. 5F**). We also noted that some of the previously identified secretome dysregulations in PMS iNSC CM were not rescued by SV treatment (e.g., IGF and inflammatory signaling). Lastly, we observed that the induction of a novel secretory signature in PMS+SV iNSC CM pertained to the regulation of neurite dynamics, glucose metabolism, and in cholesterol biosynthesis and esterification. Taken together, these results suggest that metabolic intervention via HMGCR inhibition modulates specific aspects of the PMS iNSC SASP, which pertain to the regulation of neurite dynamics and cytoprotective mechanisms.

We hypothesized that increased DSPs related to neurite dynamics in the CM of PMS iNSCs would induce a response on differentiated neurons. To functionally validate these data, we exposed SH-SY5Y neurons to the CM of Ctrl or PMS iNSCs for 72 hours (**Fig. 5G**). We found that treatment of SH-SY5Y neurons to the SASP of PMS iNSCs induced neurite retraction in comparison to CM of Ctrl iNSCs (**Fig. 5H**). Treatment of the PMS iNSCs with SV for 48 hours prevented neurite retraction of the SH-SY5Y neurons exposed to CM (**Fig. 5G-H**). Due to the significant decrease in cytoprotective factors in the SASP of PMS iNSCs, we investigated neuronal apoptosis after CM treatment by measuring cleaved caspase 3 (cl-Cas3) expression. Here we found that the SASP of PMS iNSCs induced neuronal apoptosis after 72 hours of CM exposure compared to CM of Ctrl iNSCs (**Fig. 5I**). These neurotoxic effects on the SH-SY5Y neurons were prevented by treatment of the PMS iNSCs with SV for 48 hours (**Fig. 5G,I**). Altogether our findings suggest that inhibition of HMGCR by SV treatment prevents the neurotoxic SASP in PMS iNSCs.

Overall, our work identifies a key role for cholesterol synthesis in controlling a disease-associated, lipid-related, hypermetabolic phenotype of PMS iNSCs that leads to neurotoxic signaling and is rescuable.

## DISCUSSION

Adult NSCs have been identified in demyelinated lesions in both mouse models of MS and in human tissue.^55,56^ Further analysis of NSCs in PMS lesions has found that they express markers of senescence, which was recapitulated in iPSC-NSCs generated from people with PMS.^11^ However, the role of senescent NSCs in the progression and pathology of PMS remains unclear.

Despite the importance of aging in driving neurodegenerative diseases, iPSC technologies are still predominantly employed to study cellular mechanisms of disease. Full reprogramming has been found to completely wipe the epigenetic clock back to fetal age.^29^ Conversely, direct induction of patient fibroblasts to neurons was found to preserve epigenetic and aging hallmarks^57–59^, which has also led to the development of disease modeling in a dish methodologies for neurodegenerative diseases. Here we took advantage of direct reprogramming technology for a novel patient stem cell-derived model of MS, and generated directly reprogrammed iNSCs from people with PMS, maintaining epigenetic hallmarks from the somatic cells.^28^

We identified upregulation of known senescence genes in the iPS-NSCs (ectodermal progeny) and iNSCs derived from people with PMS when compared to control counterparts, in line with previously published work.^11,60^ Interestingly no differences in senescence gene marker expression were found in somatic starting fibroblasts, or when PMS iPSCs were differentiated towards endodermal and mesodermal progenies. These results support GWAS data proposing an effect of genetic variants in genes associated with cellular senescence, mitochondrial function, and synaptic plasticity in glial cells that account for MS severity.^13,14^

Our current findings suggest that PMS iNSCs sustain a hypermetabolic state, associated with increased glycolysis and oxidative phosphorylation (OXPHOS). Similar results have been reported in erythrocytes^21^, fibroblasts^22,61^, and astrocytes^17^ of MS patients and in senescent cells^23^, without the identification of a clear function of this metabolic state in a pathophysiological context yet.

Here we used a multi-omics informed approach coupled with stable isotopic metabolic tracing to uncover a link between hypermetabolism, increased lipid synthesis, and neurotoxic protein secretion in the context of intrinsic senescence in PMS iNSCs.

We provide evidence of a direct involvement of LD accumulation and composition in reshaping saturation of the lipidome by modulating unsaturated FA trafficking between the TGs and phospholipid pools in PMS iNSCs. Our results are in line with a growing body of evidence linking hypermetabolism to significant accumulation of LDs both in the context of senescence^26^ as well as under physiological conditions in proliferative rodent NSCs.^25^ Similar findings have been reported in the context of lipid membrane restructuring in cellular stress responses.^62^

We attributed the LD accumulation in PMS iNSCs to an increased production of cholesterol and unsaturated FAs, leading to a high unsaturated ChE content. Similar lipidomic signatures, including changes in cholesterols and FAs, are being used as biomarkers of disease activity and progression in PMS.^63,64^ More specifically, elevated levels of circulating cholesterol in the CSF have been associated with adverse clinical and MRI outcomes in MS, suggesting a link between cholesterol synthesis and neuroinflammation.^65^ It is however unknown whether increased circulating cholesterol is only due to myelin destruction or is a by-product generated from CNS-infiltrating or -resident inflammatory cells. Biological changes in lipid species have also been broadly identified as a shared hallmark of neurodegenerative diseases, aging, and inflammatory conditions, such as viral infection.^66^ However, the mechanisms underpinning these changes as well as their pathophysiological implications are incompletely understood.

Our findings highlight an intrinsic, cell-autonomous, disease phenotype which emerges when subjecting a PMS genetic background to the epigenetic constraints of a NSC identity (as in Park et al.).^28^

Additionally, we uncover a dichotomic effect of SV treatment in relation to the basal metabolic state of iNSCs, incurring a high phenotypic response of lipid-laden PMS iNSCs and a comparative insensitivity of controls. Accordingly, SV may in turn be targeting, and metabolically reprogramming resident cells with high lipid anabolism in the CNS, which may include early glia such as NSC/progenitor cells, along with astrocytes and microglia. SV, which is a CNS-penetrant drug, has gone through a phase-II clinical trial (MS-STAT) for PMS, showing attenuated brain atrophy and disease progression in patients, through a yet unclear mechanism.^67^ A recent investigation employed causal models to elucidate the mechanism of action of SV in the context of PMS, drawing upon the outcomes of MS-STAT, and found the beneficial effects of SV to be independent of its effect on lowering peripheral cholesterol levels.^68^

Mechanistically, we performed multi-omics integration to provide evidence for an overarching regulatory mechanism linking the lipogenic state of PMS iNSCs and their SASP and subsequently inhibited cholesterol synthesis with SV to target the aberrant secretome. We identified a secreted signature transcriptionally co-regulated by cholesterol-dependent TFs. The activity of several of the top identified TFs that regulate the aberrant SASP of PMS iNSCs is known to be metabolically modulated by processes targeted by SV treatment in PMS iNSCs: the cholesterol metabolite itself^47–49^, the metabolic pathways of cholesterol synthesis^50^, cholesterol esterification^51^, and FA saturation.^52^ While statin-mediated modulation of senescent gene expression has been previously described^54^, SV treatment only induced a minor increase in CDKN2A expression. Therefore, we conclude that the effects of SV on the PMS iNSC-associated SASP are independent of a direct modulatory effect on the cells’ intrinsic senescent state. These results highlight the intricate complexities and heterogeneity of the ‘senescent phenotype’ and the need to reassess the defining traits of this broadly used term in a context-dependent manner.

We next sought to understand how the lipogenic state of PMS iNSCs may be mediating and leading to neurodegenerative events. Interrogating the CM, we found that the secretome of PMS iNSCs induced neuronal death and neurite retraction, while SV treatment re-wired the secretome closer to that of control iNSCs, thus rescuing the neuronal phenotype induced by PMS iNSC CM. The exact PMS iNSC secreted proteins wholly responsible for neuron cell death and retraction remain to be determined, however it is unlikely that a single candidate is responsible, while instead this is a collective effort.

Pathway analysis of the DSPs identified those related with EPH-ephrin signaling and ECM degradation, which were corrected upon SV treatment. In line with our findings, MS lesions have been found to express increased ephrin ligands and receptors, along with increased ECM molecules, believed to take part in the chronic neuroinflammation and neurodegeneration found in MS.^40,69^ Particularly, EphA4, one of the top candidates identified in our proteomic characterization as a potential mediator of PMS iNSC CM neurotoxicity, has also been implicated in driving neurodegeneration in a plethora of neurological disorders such as Alzheimer’s disease^70^, Parkinson’s disease^71^, amyotrophic lateral sclerosis, and spinal muscular atrophy.^72^ This PMS NSC secretory phenotype may therefore mediate inflammatory and neurodegenerative events by inducing reactive responses in other non-neuronal cell types prevalent in the MS brain, which were not explored in this study.

Overall, our data support the use of direct reprogramming technology to uncover new aging-associated cellular mechanisms of disease in PMS. We identified a lipogenic state that transcriptionally promotes neurotoxic SASP leading to neurite retraction and neuronal death. We further uncovered how SV/HMGCR inhibition rescues this state via metabolic reprogramming and restructuring of the lipidome and secretome. In addition, we have generated large omics-based repositories, including metabolomics, proteomics, and lipidomics datasets, which can serve for further mining by the MS research community. Additional studies are needed to uncover why PMS cells have intrinsic metabolic dysfunction and if it is specifically linked to senescence or is a unique phenotype of PMS.

By uncovering inherent dysfunction within stem cells derived from patients, our research also lays pivotal groundwork for critically assessing autologous stem cell therapies in MS. This underscores the potential for these patient-specific cells to carry distinct disease traits and internal dysfunctions, urging a reconsideration of their use in treatment approaches.^73^

## LIMITATIONS OF STUDY

Compared to most stem cell-based disease modelling approaches for monogenic diseases, our study lacks a classical direct control given by isogenic lines, something that is at present not feasible for MS research. Given that we generated cell lines from individuals with possibly different genetic backgrounds, our study may also be falling short in study power to identify more subtle yet pathologically relevant features. While being very systematic in the characterization of paracrine mechanisms of cellular toxicity and injury, our work is not addressing the yet relevant question of whether the identified disease associated dysfunctional cellular phenotype may also affect cell-to-cell interactions. Hence, multicellular 2D and/or 3D cultures refined around the study of interactions between NSCs and neuronal *vs*. non neuronal cells hold the promise to further understand the role and function of disease associated NSCs in PMS.

## ACKNOWLEDGMENTS

The authors wish to acknowledge L. Ionescu, V. Pappa, G. Pluchino, O. Hruba, S. Rizzi, D. Sastre-Stanescu, M. Sciacovelli, A. Speed, A. Tolkovsky, D. Nerguizian, and F. Gamboni for technical and intellectual inputs throughout this study. The authors also thank M. Whitehead, for providing the neuronal SH-SY5Y cells.

This research was supported by the Ferblanc Foundation G112716 (SP and AMN); National MS Society Research Grant RG 1802-30200 (SP and LPJ); Bascule Charitable Trust RG98181 (SP); Wings for Life RG 82921 (SP and LPJ), and the Fondazione Italiana Sclerosi Multipla FIMS 2018/R/14 (SP and LPJ) and 2022/R-Single/011 (SP). RBI is supported through an MRC-DTP and Cambridge Trust studentship and consumable award (RG86932) and St. Edmund’s College Tutorial Award. AMN is the recipient of a European Committee for Treatment and Research in Multiple Sclerosis (ECTRIMS) Postdoctoral Research Fellowship Exchange Program fellowship (G104956) and is supported through a UK MS Society Centre Excellence grant (G118541). ECW is supported through an MRC-DTP iCASE PhD studentship award, jointly funded by AstraZeneca, (G117817). IM is supported by Wellcome Trust (203151/Z/16/Z) and the UKRI Medical Research Council (MC_PC_17230). PP is supported through an MRC-DTP and Cambridge Trust PhD studentship and consumable award (RG86932) and Queen’s College Tutorial Award. LPJ was supported by a Fondazione Italiana Sclerosi Multipla FIMS and Italian Multiple Sclerosis Association AISM Senior research fellowship financed or co financed with the ‘5 per mille’ public funding cod. 2017/B/5 (LPJ), a Wellcome Trust Clinical Research Career Development Fellowship (G105713), and a National MS Society Research Grant RFA-2203-39318. CMW is the recipient of a National MS Society (USA) Post-doctoral fellowship (FG-2008-36954). FE is supported through the Austrian Science Fund (FWF) (Special Research Programme F7804-B; I 4791; TAI 801) and EJP European Joint Programme on Rare Diseases (I 5184).

## AUTHOR CONTRIBUTIONS

Conceptualization, RBI, AMN, AD, CF, and SP; Methodology, RBI, AMN, JAR, ECW, PP, MD, DS, MSC, LPJ, LP, CMW, CF, IM, AD, and SP; Investigation, RBI, AMN, JAR, ECW, PP, MD, DS, LP, CMW; Writing – Original Draft, RBI and AMN.; Writing – Review & Editing, RBI, AMN, JAR, ECW, LPJ, IM, AD, and SP.; Funding Acquisition, RBI, AMN, PP, LPJ, CMW, AD, IM, and SP; Resources, MSC, VF, FE, and TL; Supervision, CF, IM, AD, and SP.

## METHODS

### Lead contact

Further information and requests for resources and reagents should be directed to and will be fulfilled by the lead contact Stefano Pluchino.

### Materials availability

This study did not generate new unique reagents.

### Data and code availability

- All raw data to be uploaded to appropriate repositories upon acceptance of manuscript
- Uncropped westerns provided in Data S1
- Raw values from graphs will be provided as Data S1 upon revision of manuscript
- The original code used in this paper has been added to a github repository (https://github.com/Core-Bioinformatics/IonescuNicaise2023)
- Any additional information required to reanalyze the data reported in this paper is available from the lead contacts upon request.

### EXPERIMENTAL MODEL AND SUBJECT DETAILS DETAILS

The cohort consists of 4 PMS (3 SPMS and 1 PPMS) and 3 healthy control donors between 25 and 63 years of age. The cohort includes representation from both genders (5 male and 2 female), evenly distributed across PMS and control groups (**Table 1**). PMS donors underwent clinical assessment when recruited for the study. Fibroblasts were provided by the New York Stem Cell Foundation (NYSCF) Research Institute through their Repository (http://www.nyscf.org/repository).^74^ Patients were recruited at the Tisch Multiple Sclerosis Research Center of New York, upon informed consent and institutional review board approval (BRANY). Control fibroblasts from lines A and B (**Table 1**) were generated from adult dermal fibroblasts after obtaining consent and ethical clearance by the ethics committee of the University of Würzburg, Germany. All other fibroblasts used in this study were obtained from NYCSF.

#### Derivation and reprogramming of skin fibroblasts to iPSCs and iNSCs

Derivation and reprogramming of skin fibroblasts to iPSCs was performed as previously described for the NYSCF cohort.^75^ Briefly, 3-5 mm skin biopsies were collected in Biopsy Collection Medium (RPMI 1460 [Thermo Fisher] with 1X Antibiotic-Antimycotic [Thermo Fisher]), cut into smaller pieces (<1 mm) and plated onto a TC-treated 35 mm dish in Biopsy Plating Medium, composed by Knockout DMEM (Thermo Fisher), 2 mM GlutaMax (Thermo Fisher), 0.1 mM non-essential amino acids (Thermo Fisher), 0.1 mM β-Mercaptoethanol (Thermo Fisher), 10% Fetal Bovine Serum (FBS) (Thermo Fisher), 1X Penicillin-Streptomycin (P/S) (Thermo Fisher) and 1% Nucleosides (Millipore). Once the first fibroblasts migrated out of the biopsies, the cultures were maintained in growth medium (DMEM Glutamax I [Thermo Fisher] supplemented with 10% fetal bovine serum, 1% non-essential amino acids and 1 mM sodium pyruvate (Thermo Fisher) at 37°C with 5 % CO_2_ and fed every 3-4 days. After reaching 90% confluency the fibroblasts were detached with trypsin-EDTA 0.05% for 5 min followed by neutralization in DMEM and spun down at 300xg for 5 min. They were split 1:4 into growth media.

Fibroblasts at passage 3-5 were reprogrammed to iPSCs using the integration free technology based on non-modified RNA plus microRNA (kit from REPROCELL, formerly Stemgent), following manufacturer’s instructions. About 25×10^3^ fibroblasts/well were plated onto Matrigel-coated 12-well plates in culture medium for 24 hours and then in NuFF-conditioned Pluriton reprogramming medium with B18R. Cells were transfected for 11 consecutive days using Stemfect as following: day 0 microRNA only, days 1-3 RNA only, day 4 microRNA plus RNA, days 5-11 RNA only. From day 11, TRA-1-60^+^ colonies (live stained) were manually picked and re-plated on mouse embryonic fibroblasts in HUESM medium (Knockout-DMEM, 20% knock-out serum, glutamax 2mM, NEAA 0.1mM, 1X P/S and β-mercaptoethanol 0.1mM [Thermo Fisher]). Pluripotent colonies were passaged and adapted to feeder-free conditions with hESC matrigel (Corning) and mTeSR1 (STEMCELL Technologies) medium. The WIBJ iPSC line, obtained from Cambridge BioResource as a gift from Alessandra Granada, was reprogrammed from fibroblasts using the CytoTune 1 non-integrating kit (Thermo Fisher). iPSC media was changed every day. When confluent, cells were lifted using accutase (Thermo Fisher), spun at 300xg for 5 min, and split 1:10-1:20 onto hESC-matrigel coated plates with Y-27632 (10 uM) (Thermo Fisher) in mTeSR1 media.

To generate directly induced neural stem cells (iNSCs) from fibroblasts we used a nonintegrating Sendai virus-based direct conversion strategy, as previously described.^27^ Briefly, fibroblasts were seeded at 75,000/well in fibroblast media in a non-coated 12-well. On the next day the fibroblasts were transduced using the CytoTune-iPS 2.0 Sendai Reprogramming Kit (Thermo Fisher) with hKOS (MOI: 3), hc-Myc (MOI: 3), hKlf4 (MOI: 3). The day following transfection, the medium was switched to neural induction medium (NIM) [DMEM:F12 and Neurobasal (1:1), supplemented with N2 supplement (1x, ThermoFisher), 1% glutamax, B27 supplement (1x, ThermoFisher), CHIR99021 (3 µM, Cell Guidance Systems), SB-431542 (2 µM, Cayman Chemicals) and hLIF (10 ng/ml, Cell Signaling Technology)], and cells were moved to 39°C with 5% CO_2_ to achieve viral clearance over 14 days. A few samples were collected here to generate positive controls for quality control assays. Medium changes were performed every other day. Following 25 days of transfection, iNSC colonies were manually selected, seeded onto Growth Factor Reduced (GFR) Matrigel Matrix (1:20 in DMEM/F12) coated plates for expansion, and subjected to quality control assays. iNSCs were maintained in NIM media until 70% confluent, then lifted using accutase (ThermoFisher), spun at 300xg for 3 mins, and plated onto GFR-Matrigel coated plates with Y-27632 (10 µM, Miltenyi Biotec) between 1:3-1:5 in NIM media. Media was changed every second day as needed. Experiments were performed on cells from passages 20-40.

### METHOD DETAILS

#### Trilineage Differentiation

Trilineage differentiation of the iPSCs was achieved using the STEMdiff Trilineage Differentiation Kit (STEMCELL Technologies). Mediums were prepared based on kit instructions. For ectodermal differentiation iPSCs were plated at 200,000 cells/cm^2^, mesoderm at 50,000 cells/cm^2^, and endoderm 200,000 cells/cm^2^. Mesoderm and endodermal differentiations were fed with indicated mediums every day for 5 days, then collected for RNA isolation or fixed for immunocytochemistry. Ectodermal differentiations were fed every day for 7 days then collected.

#### RNA Isolation, PCR, and qRT-PCR

Total RNA from all cell types was isolated using the RNeasy Mini Kit (Qiagen) following manufacturer’s instructions. Briefly, cells were washed with ice-cold PBS followed by the addition of RLT lysis buffer. Samples were then stored at −80°C until extraction. Total RNA was then quantified with the NanoDrop 2000c instrument (Thermo Fisher).

iNSC quality control was performed using PCR analysis. Here 300 ng of RNA was reverse-transcribed using the High Capacity cDNA Reverse Transcription Kit (Thermo Fisher) according to the manufacturer’s instructions using a 20 uL reaction volume on a T100 Thermal Cycler (BioRad, 1861096). The RT-PCR reaction was made using DreamTaq buffer (Thermo Fisher), dNTPs (2 mM each, Thermo Fisher), forward and reverse primers (0.5 uM, Table S2), DreamTaq Hot Start DNA Polymerase (Thermo Fisher), and finished to 24 uL with water. 1 uL of cDNA from each sample was loaded into the PCR reaction. Oct4 reactions were cycled at: 95°C for 3 mins; x30 cycles of 95°C 30 sec, 60°C 30 sec, 72°C 1 min; 72°C 5 min, hold at 4°C. SeV, KOS, cMyc, SOX2, Pax6, and nestin reactions were cycled at: 95°C for 3 mins; x30 cycles of 95°C 30 sec, 55°C 30 sec, 72°C 1 min; 72°C 5 min, hold at 4°C. Samples were diluted with Gel Loading Dye (New England BioLabs), and 5 μL was loaded into a 2% agarose gel (Thermo Fisher) in 1X TAE buffer (MP Biomedicals). TrackIt 100 bp DNA Ladder (Thermo Fisher) was used to assess band size. DNA was visualized using Gelred Stain (Biotium) in the gel and imaged on a BioRad XR GelDoc Imager.

For qRT-PCR analysis, 500 ng of RNA was reverse-transcribed using the High Capacity cDNA Reverse Transcription Kit (Thermo Fisher) according to the manufacturer’s instructions using a 20 μL reaction volume on a T100 Thermal Cycler (BioRad). qRT-PCR was performed with the TaqMan Fast Universal PCR Master Mix (2x) (Thermo Fisher) and TaqMan Gene Expression Assays (see resources table). Ribosomal subunit 18S was used to normalize gene expression. 1.5 μL of cDNA from each sample was run in duplicate using a QuantStudio 7 Flex (Thermo Fisher) and analyzed with the 2^-ΔΔCT^ method.

#### Immunocytochemistry for iNSCs, iPSCs, and Trilineage Differentiation

Cells were plated at a density between 40,000 – 60,000 cells/cm^2^ on GFR-matrigel coated glass coverslips. Cells were fixed for 10 minutes with 4% paraformaldehyde (Thermo Fisher) then permeabilized for 10 minutes in PBS with 0.25% Triton-X100 (Thermo Fisher). Cells were then blocked for 1 hour using 1% normal donkey serum (Thermo Fisher) in PBS with 0.1% Triton-X100. Cells were incubated with the following primary antibodies in the above blocking buffer: Nestin (Novus Biolgicals, 1:500), ETNPPL (Atlas, 1:500), SOX2 (Abcam, 1:500), SSEA4 (Thermo Fisher, 1:100), Nanog (Abcam, 1:1000), Oct4 (Santa Cruz, 1:200), SOX17 (R&D, 1:200), ACTA2 (antibodies.com, 1:20), Pax6 (BioLegend, 1:100), SOX1 (R&D, 1:50), Oct4 (Reprocell, 1:250), TRA-1-60 (EMD Millipore, 1:250), TRA-1-81 (EMD Millipore, 1:250), NANOG (Cell Signaling, 1:100), SSEA4 (Abcam, 1:250), and SOX2 (Reprocell, 1:250) at 4°C overnight. Coverslips were washed three times for 10 minutes each in PBS with 0.1% Triton-X100 then incubated with species appropriate secondary antibodies (1:1000) in blocking buffer for one hour at room temperature. Coverslips were washed again as above then stained with DAPI in PBS (300 nM, Thermo Fisher) for five minutes then washed in PBS. Coverslips were mounted onto glass slides using Fluoromount-G (Thermo Fisher). Images were taken on a Leica DMI400B microscope at 40X in oil immersion.

#### Immunoblotting

Cells were homogenized in 1x RIPA buffer (Abcam) supplemented with 1x protease and phosphatase inhibitors (Thermo Fisher). Equal protein amounts (10 µg) were resolved by SDS-PAGE on 4-15% Mini-PROTEAN TGX Stain Free gels (30 μL, 10 well comb, BioRad, Cat# 4568083) and transferred to polyvinylidene fluoride (PVDF) membranes using a Trans-Blot Turbo Transfer Pack (BioRad, Cat# 1704156). Membranes were blocked with 5% milk in 0.1% TBST for one hour at room temperature. Membranes were then immunoblotted over night at 4°C with the indicated primary antibodies diluted in blocking buffer: Phosphorylated Stat1 (Tyr701) (D4A7) (Cell Signaling, 1:1000), STAT1 (Cell Signaling, 1:1000), ISG15 (Cell Signaling, 1:1000), β-actin (Thermo Fisher, 1:5000). Membranes were washed three times with TBST for 5 minutes and immunoblotted with the appropriate HRP-conjugated secondary antibodies: anti-rabbit (Cell Signaling, Cat#7074, 1:2000) or anti-mouse (Cell Signaling, Cat#7076, 1:2000). Membranes were washed three times with TBST for 5 minutes, then imaged on a ChemiDoc MP Imaging System (BioRad), soaked in Clarity Western ECL Substrate (Cat#, 1705061). Protein band densitometry was measured in ImageJ. An identical selection frame was used to determine the mean gray value (MGV) measurement of protein bands and background areas above and below each band. Band intensity (BI) was depermined by the formula: BI = (255-MGV_protein band_) − [(255 − MGV_background above_) + (255 − MGV_background below_)]/2. BI values for PMS iNSC lines were normalized to the BI values of control iNSC lines within each blot.

#### Immunocytochemistry for mitochondrial morphology assay

Cell lines were plated in technical triplicates at a density between 40,000 – 60,000 cells/cm^2^ on GFR-matrigel coated glass coverslips. Cells were fixed for 10 minutes with 4% paraformaldehyde then washed twice with PBS for 5 minutes. Cells were then blocked for 1 hour using 1% normal goat serum in PBS with 0.1% Triton-X100. Cells were incubated with TOMM20 (Abcam, 1:1000) at 4°C overnight in the above blocking buffer. Coverslips were washed again as above then stained with DAPI in PBS (300 nM) for five minutes then washed in PBS. Coverslips were mounted onto glass slides using Fluoromount-G. 4 regions of interest (ROIs) were imaged on each coverslip, acquiring 36 Z-stacks with a step size of 0.2 µm on a Leica SPE Confocal microscope at 63X in oil immersion and analyzed using Fiji. Maximal projection images of each ROI were generated for the TOMM20 channel, and 48 random single-cell selections were made (4 cells/ROI, 3 ROIs/coverslip, 3 coverslips/cell line). The resulting single-cell selections were analyzed using the MiNA plugin.^76^ MiNA estimates mitochondrial footprint from a binarized copy of the image as well as the lengths of mitochondrial structures using a topological skeleton.

#### Seahorse mitochondrial stress test

A Seahorse calibration plate was rehydrated with XF calibration solution (Agilent) overnight in a non-CO_2_ incubator at 37°C. On the same day, fibroblasts were seeded at 45 × 10^3^ cells per well (5-10 replicates per cell line) in fibroblast growth media in a Seahorse XFe24 Microplate (Agilent, 100777-004) and incubated for 24 hours to allow cells to adhere. Similarly, iNSCs were seeded at 12 × 10^4^ cells per well (5-10 replicates per cell line) in NIM media in a Seahorse XFe24 Microplate coated with GFR-matrigel and incubated for 24 hours to allow cells to adhere. In each microplate 4 wells were left empty to serve as background wells, as per manufacturer’s protocol guidelines. The calibration plate was then calibrated using the Seahorse instrument. The cells were washed once with DMEM Assay Medium (XF DMEM [Agilent] supplemented with XF glucose [10 mM, Agilent], XF glutamine [2 mM, Agilent] and XF pyruvate [1 mM, Agilent]), then left in DMEM Assay Medium and placed in a non-CO_2_ incubator for 1 hour. Cells were then subjected to the XFe Cell Mito stress test (Agilent) on a Seahorse XFe96 Analyzer (Agilent) according to the manufacturer’s instructions. The cell plate was equilibrated, and the baseline was measured. Oligomycin was used to inhibit mitochondrial ATP production (2 µM final concentration in well). FCCP was used to induce mitochondrial uncoupling and maximal mitochondrial respiration (2 µM final concentration in well). Rotenone/antimycin A were used to inhibit mitochondrial complex I and complex III respectively, and therefore stop mitochondrial respiration (1 µM final concentration in well). Three reads were obtained (every 8.75 minutes) at baseline and after each drug injection. If cells did not respond to the addition of drugs, they were excluded from further analysis. After the mitochondrial stress test on fibroblasts protein levels of each well were measured from the cell lysates by Pierce BCA Protein Assay and used for signal normalization. Due to the use of extracellular matrix coating for iNSC culturing, normalization by nuclear count was performed. After the mitochondrial stress test iNSC plates were fixed with 4% paraformaldehyde, stained with DAPI and 9 ROIs were captured per well at 10X magnification on a Leica DMI400B microscope. Nuclear counts were determined using the Threshold, Watershed, and Analyze particles functions in Fiji and used for signal normalization. Analysis was performed using the Wave 2.6.1. software (Agilent).

#### Cell size, mitochondrial mass, and membrane potential

iNSCs were plated in technical duplicate, in two separate experiments, at a density of 800,000cells/well on GFR-matrigel coated 6-well plates. Cells were lifted with accutase, spun for 5minutes at 300 × g and washed once with 1x staining buffer (BioLegend). iNSCs were stained in suspension for 15 minutes at 37°C on an orbital shaker (500 rpm), with 100 nM of the mitochondrial mass indicator MitoTracker Green (Invitrogen), 50 mM of the mitochondrial membrane potential indicator (MitoTracker Red CMXRos FM) and 1x of the live/dead stain Zombie Violet diluted in staining buffer. Cells were washed once with staining buffer and analyzed by flow cytometry on a BD LSRFortessa Flow Cytometer (BD Biosciences), recording 100,000 total events. Analysis was performed using FlowJo v10 (BD Biosciences). First a single cell population was gated using FSC-H by FSC-A, next live cells were gated using the ZombieViolet negative population. The mean cell size of iNSCs was estimated from the geometric mean of the forward-scatter area (FSC-A) parameter in the gated population. Histograms of red (mitochondrial membrane potential) and green (mitochondrial mass) fluorescence were assessed, and geometric means (i.e., mean fluorescence intensities) were extracted. The mitochondrial mass signal was used for normalization of the mitochondrial membrane potential signal. Resulting values for PMS iNSCs were further normalized to control levels set to a value of 1.

#### Mitochondrial superoxide levels

iNSCs were plated in technical duplicate, in two separate experiments, at a density 800,000 cells/well on GFR-matrigel coated 6-well plates. Cells were lifted with accutase, spun for 5 minutes at 300 × g and washed once with 1x staining buffer (BioLegend). iNSCs were stained in suspension for 30 minutes at 37°C on an orbital shaker (500 rpm), using 5 µM of the mitochondrial superoxide indicator MitoSox (Invitrogen) and 1x of the live/dead stain Zombie Violet diluted in staining buffer. Cells were washed once with staining buffer and analyzed by flow cytometry on a BD LSRFortessa Flow Cytometer (BD Biosciences), recording 100,000 total events. Analysis was performed using FlowJo v10 (BD Biosciences). First a single-cell population was gated using FSC-H by FSC-A, next live cells were gating using the ZombieViolet negative population. The histogram of red fluorescence was assessed, and the geometric mean (i.e., mean fluorescence intensity) was extracted. Resulting values for PMS iNSCs were further normalized to control levels set to a value of 1.

#### Mitochondrial copy number assay

Total DNA from all cell types was isolated using the DNeasy Blood & Tissue Kit (Qiagen) according to manufacturer’s instructions. Total DNA was then quantified with the NanoDrop 2000c instrument (Thermo Fisher). For each sample, 20 ng of DNA was loaded into a qPCR reaction containing the indicated primers for MT-ND2 and the TaqMan Fast Universal PCR Master Mix (Thermo Fisher) and ran on a QuantStudio 7 Flex (Thermo Fisher). The RNase P TaqMan Copy Number Reference Assay was used for normalization within the same sample, using the with the 2^-ΔΔCT^ method.

#### Immunocytochemistry for lipid droplet quantification

iNSCs were plated in technical triplicates at a density between 40,000 – 60,000 cells/cm^2^ on GFR-matrigel coated 13 mm diameter glass coverslips. Fibroblasts were plated in technical triplicates at a density between 25,000 – 37,500 cells/cm^2^ on GFR-matrigel coated 13 mm diameter glass coverslips. Cells were fixed for 10 minutes with 4% paraformaldehyde then washed twice with PBS for 5 minutes. Lipid droplets were stained using the 1x LipidSpot610 dye for 20 minutes at room temperature. Coverslips were then counter stained with DAPI in PBS (300 nM) for five minutes then washed in PBS and mounted onto glass slides using Fluoromount-G (Thermo Fisher). 3 ROIs were imaged on each coverslip using a Leica DMI400B microscope at 63X in oil immersion and batch analyzed using Fiji. Lipid droplet numbers and nuclei numbers were measured using the Threshold, Watershed, and Analyze particles functions in Fiji, and lipid droplet content was calculated as a ratio of lipid droplet count/nuclei count.

#### High resolution imaging and 3D reconstruction of lipid droplets

Cell lines were plated at a density between 40,000 – 60,000 cells/cm^2^ on GFR-matrigel coated 8-well glass chamber slides. Cells were fixed for 10 minutes with 4% paraformaldehyde. Plasma membrane was stained using BioTracker 555 Orange Cytoplasmic Membrane Dye (1:1000) and lipid droplets were stained with 1x LipidSpot™ in PBS for 20 minutes at room temperature. Nuclei were stained with 300 nM DAPI in 1x PBS. Cells were then washed with 1x PBS twice for 5 minutes. Z-stack Lightning imaging with a 0.2 µm step size was performed using confocal microscope Leica Stellaris8 on a 63x objective and oil immersion setting. 3D surfaces for each staining where then rendered using the Imaris 9.5 software.

#### Sample generation for metabolomics, stable isotope tracing, proteomics, and lipidomics

iNSCs were seeded in NIM with Y-27632 (10 µM) at a density of 100,000 cells/cm^2^ GFR-matrigel coated wells, in technical triplicates per line. Medium was changed the next day. Glucose tracer experiments were run on day 2 after seeding by removing the culture medium and adding glucose-free NIM [DMEM:F12 w/o glucose and Neurobasal (1:1), supplemented with N2 supplement (1x, ThermoFisher), Glutamax (Thermo Fisher, 3.2 mM), pyruvate (Agilent, 0.35 mM), B27 supplement (1x, ThermoFisher), CHIR99021 (3 µM, Cell Guidance Systems), SB-431542 (2 µM, Cayman Chemicals) and hLIF (10 ng/ml, Cell Signaling Technology)] supplemented with D-Glucose-^13^C_6_ (ThermoFisher, 20.4 mM). Samples were collected after 30 minutes (for optimal resolution of glycolytic pathway), 6 hours (for assessment of protein acetylation patterns) and 24 hours (for optimal resolution into TCA cycle and anabolic metabolic pathways). The glutamine tracer experiment was run on day 2 after seeding by removing the culture medium and adding glutamine-free NIM [DMEM:F12 w/o glutamine and Neurobasal (1:1), supplemented with N2 supplement (1x, ThermoFisher), glucose (Agilent, 20.4 mM), pyruvate (Agilent, 0.35 mM), B27 supplement (1x, ThermoFisher), CHIR99021 (3 µM, Cell Guidance Systems), SB-431542 (2 µM, Cayman Chemicals) and hLIF (10 ng/ml, Cell Signaling Technology)] supplemented with L-Glutamine-^13^C_5_,^15^N_2_ (Thermo Fisher, 3.2 mM). Samples were collected after 24 hours (for optimal resolution into TCA cycle and anabolic metabolic pathways). Samples for intracellular and extracellular lipidomics analysis were collected on day 3 after seeding (no other medium changes were done except for the day 1 feeding with fresh NIM). Sample collection was performed on ice. Conditioned culture medium was collected, spun at 300 × g for 5 minutes to remove cellular debris and 100 µL was stored at −80 °C until sample processing. Cells were then lifted using accutase and spun at 300 × g for 5 mins. Cell pellets were resuspended in PBS for counting and spun once more at 300 × g for 5 mins. Supernatants were aspirated and cell pellets were stored at −80°C until sample processing. Mass spectrometry-based omics analyses were performed on frozen cell pellets at the University of Colorado School of Medicine Mass Spectrometry Facility, a shared resource of the University of Colorado Cancer Center.

#### Sample processing for metabolomics and stable isotope tracing

Metabolites from frozen pellets were extracted at 2e6 cells per mL using ice cold 5:3:2 methanol:acetonitrile:water (v/v/v) with vigorous vortexing at 4 °C followed by centrifugation as described. Clarified supernatants were analyzed (10 µL per injection) by ultra-high-pressure liquid chromatography coupled to mass spectrometry on a Vanquish UHPLC (Thermo Fisher) coupled to a Q Exactive mass spectrometer (Thermo Fisher) in positive and negative ion modes (separate runs). The UHPLC utilized a 5 min C18 gradient at 450 µL/min; eluate was introduced to the MS via electrospray ionization as previously described in detail. Data analysis and quality control measures were performed as described. Isotopologue data from stable isotope tracing experiments was corrected for natural abundance then plotted using GraphPad Prism 10.0.

#### Sample processing for proteomics analysis

The protein pellets after metabolomics extraction were solubilized in 4% SDS in 100 mM triethylammonium bicarbonate (TEAB) pH 7.0 lysis buffer. The samples were digested in the S-Trap micro spin column (Protifi, Huntington, NY) following the manufacturer’s procedure. Samples were reduced with 10 mM DTT at 55°C for 30 min, cooled to room temperature, and then alkylated with 25 mM iodoacetamide in the dark for 30 min. Next, a final concentration of 1.2% phosphoric acid and then six volumes of binding buffer (90% methanol; 100 mM triethylammonium bicarbonate, TEAB; pH 7.0) were added to each sample. After gentle mixing, the protein solution was loaded to a S-Trap micro spin column, spun at 1500 × g for 2 min, and the flow-through collected and reloaded onto the S-Trap micro spin column. This step was repeated three times, and then the S-Trap micro spin column was washed with 400 μL of binding buffer 3 times. Finally, 1 μg of sequencing-grade trypsin (Promega) and 125 μL of digestion buffer (50 mM TEAB) were added onto the filter and digested carried out at 37 °C for 6 hours. To elute peptides, three stepwise buffers were applied, with 100 μL of each with one more repeat, including 50 mM TEAB, 0.2% formic acid in water, 50% acetonitrile, and 0.2% formic acid in water. The peptide solutions were pooled, lyophilized, and resuspended in 100 μl of 0.1% FA.

Conditioned media samples were digested according to the FASP protocol using a 10 kDa molecular weight cutoff filter. In brief, samples were mixed in the filter unit with 8 M urea, 0.1 M ammonium bicarbonate (AB) pH 8.0, and centrifuged at 14,000 g for 15 min. The proteins were reduced with 10 mM DTT for 30 min at RT, centrifuged, and alkylated with 55 mM iodoacetamide for 30 min at RT in the dark. Following centrifugation, samples were washed 3× with 8 M urea, and 3× with 50 mM AB, pH 8.0. Protein digestion was carried out with sequencing grade modified trypsin (Promega) at 1/50 protease/protein (wt/wt) at 37°C overnight. Peptides were recovered from the filter using 50 mM AB. Samples were dried via Speed-Vac and desalted and concentrated on Thermo Scientific Pierce C18 Tip.

#### Proteomics data acquisition

A 20 μL volume of each sample was loaded onto individual Evotips for desalting and then washed with 20 μL 0.1% FA followed by the addition of 100 μL storage solvent (0.1% FA) to keep the Evotips wet until analysis. The Evosep One system (Evosep, Odense, Denmark) was used to separate peptides on a Pepsep column, (150 um inter diameter, 15 cm) packed with ReproSil C18 1.9 μm, 120 A resin. The system was coupled to the timsTOF Pro mass spectrometer (Bruker Daltonics, Bremen, Germany) via the nano-electrospray ion source (Captive Spray, Bruker Daltonics).

The mass spectrometer was operated in PASEF mode. The ramp time was set to 100 ms and 10 PASEF MS/MS scans per topN acquisition cycle were acquired. MS and MS/MS spectra were recorded from m/z 100 to 1700. The ion mobility was scanned from 0.7 to 1.50 Vs/cm^2^. Precursors for data-dependent acquisition were isolated within ± 1 Th and fragmented with an ion mobility-dependent collision energy, which was linearly increased from 20 to 59 eV in positive mode. Low-abundance precursor ions with an intensity above a threshold of 500 counts but below a target value of 20000 counts were repeatedly scheduled and otherwise dynamically excluded for 0.4 min.

#### Proteomics Database Searching and Protein Identification - *Cells*

MS/MS spectra were extracted from raw data files and converted into .mgf files using MS Convert (ProteoWizard, Ver. 3.0). Peptide spectral matching was performed with Mascot (ver. 2.5) against the Uniprot human database. Mass tolerances were +/- 15 ppm for parent ions, and +/- 0.4 Da for fragment ions. Trypsin specificity was used, allowing for 1 missed cleavage. Met oxidation, protein N-terminal acetylation, Lys C13 acetylation and peptide N-terminal pyroglutamic acid formation were set as variable modifications with Cys carbamidomethylation set as a fixed modification.

Scaffold (version 5.0, Proteome Software, Portland, OR, USA) was used to validate MS/MS based peptide and protein identifications. Peptide identifications were accepted if they could be established at greater than 95.0% probability as specified by the Peptide Prophet algorithm. Protein identifications were accepted if they could be established at greater than 99.0% probability and contained at least two identified unique peptides.

#### Database Searching and Protein Identification – *Conditioned media*

Raw data files conversion to peak lists in the MGF format, downstream identification, validation, filtering, and quantification were managed using FragPipe version 13.0. MSFragger version 3.0 was used for database searches against a mouse database with decoys and common contaminants added. The identification settings were as follows: Trypsin, Specific, with a maximum of 2 missed cleavages, up to 2 isotope errors in precursor selection allowed for, 10.0 ppm as MS^1^ and 20.0 ppm as MS^2^ tolerances; fixed modifications: Carbamidomethylation of C (+57.021464 Da), variable modifications: Oxidation of M (+15.994915 Da), hydroxylation of P (+15.994915 Da), acetylation of protein N-term (+42.010565 Da), pyrolidone from peptide N-term Q or C (−17.026549 Da).

#### Protein acetylation

The ^13^C acetylated peptides were identified using the Mascot search algorithm by searching all MS/MS spectra against the human database for [^13^C_2_]lysine as a variable modification. The quantitative MS analysis of the ^13^C acetylation sites were performed based on the spectral count data (i.e. peptide spectral matches).

#### Lipidomics sample processing

Lipids were extracted from frozen cells at 2e6 cells/mL and from conditioned media at 1:25 dilution with cold MeOH. Suspensions were briefly vortexed and placed at −20 °C for 30 min followed by a vigorous vortex for 30 min at 4 °C. Insoluble material was pelleted by centrifugation (18,213 × g, 10 min, 4 °C) and supernatants were isolated for analysis. Lipidomics analysis employed a Thermo Vanquish UHPLC system coupled to a Thermo Q Exactive mass spectrometer. 5 uL injections of the samples were resolved across a 2.1 × 30 mm, 1.7 μm Kinetex C18 column (Phenomenex) using a 5-minute, reverse-phase gradient adapted from a previous method. The Q Exactive was run independently in positive and negative ion mode, scanning using full MS from 125-1500 m/z at 70,000 resolution and top 10 data-dependent MS^2^ at 17,500 resolution. Electrospray ionization was achieved with 45 Arb sheath gas, 25 Arb auxiliary gas, and 4 kV spray voltage. Calibration was performed prior to the run using the PierceTM Positive and Negative Ion Calibration Solutions (Thermo Fisher). Run order of samples was randomized and technical replicates were injected after every 4 samples to assess quality control. Raw files were searched against lipid databases using LipidSearch (Thermo Fisher).

#### Metabolomic and lipidomic analysis

Metabolomics and lipidomics datasets were pre-processed using per-total normalization and feature standardization, on mean and standard deviation. Principal component analysis (PCA) was performed using prcomp (from stats v4.2.1 R package) function. Metabolic pathway level analysis of metabolomics data was performed by uploading separate lists of differentially abundant intracellular and extracellular metabolites to Metaboanalyst 5.0. Pathway Enrichment webtool. Pathway analysis parameters consisted of enrichment by Hypergeometric test and topology analysis by Relative-betweenness Centrality. All compounds in the *Homo sapiens* (SMPDB) pathway library were used as reference metabolome. Scatter plot was chosen as visualization method. Lipid moiety analysis for the intracellular lipidome was performed using the Substructure Analysis function of the LINEX^2^ webtool.^35^ Lipid pathway level analysis of the intracellular lipidomics data was performed using the Bioinformatics Methodology for Pathway Analysis (BioPAN) webtool.

#### Proteomics analysis

Proteomics modalities were pre-processed using minimum value imputation of missing values, followed by median normalization.^77^ Further analyses were performed using bulkAnalyseR v1.1.0^41^ with differential expression calls using edgeR v3.40.2^78^ and enrichment analysis performed on differentially expressed proteins using gprofiler2 v0.2.2^79^; the set of all genes expressed in matching RNAseq data was used as the background, only proteins expressed in >20% of cell lines were considered and FDR multiple testing correction was also performed.

#### Multi-omics factor analysis (MOFA)

Multi-omics factor analysis (MOFA) was used to integrate intracellular lipidomics and secretomics modalities, using the MOFA2 v1.8.0 R package.^53^ Replicates per cell line were pooled (by averaging); per-total normalization and feature scaling were also performed. Any proteins expressed in less than 20% of cell lines were removed from subsequent steps. MOFA imputation was performed for missing values in the secretomics data. A MOFA model was created using 3 factors (as recommended) and otherwise default settings. The calculate_variance_explained() and correlate_factors_with_covariates() functions from MOFA2 were used to assess the % variance explained by each factor, in each modality, and the correlation between each factor with disease and treatment covariates.

We noted a large number of missing values in the extracellular proteomic (secretomic) dataset, due to the detection limitations of the mass spectrometry-based method employed. In some cases, a consistent trend could be observed where only samples pertaining to a particular experimental group consistently produced a detectable signal, which was suggestive of a biologically relevant feature. In other cases, the signal across samples followed an inconsistent pattern, oscillating below and above the detection limit indiscriminate of a particular experimental group, which suggested technical variation. To prevent misattributing biological relevance to differences emerging from technical variation across samples we pre-filtered our dataset and removed any proteins expressed in less than 20% of cell lines from our analysis. The remaining missing values were imputed using the MOFA imputation approach; we acknowledge the impact the missing values and their imputation with limited information may have on conclusions.

#### Multi-omics covariation network analysis

Multi-omics covariation network inference was performed using GENIE3 v1.20.0^80^ across modalities and visualized using visNetwork v2.1.2; replicates were pooled per cell line (by averaging across replicates); all modalities were pre-processed using per-total normalization and feature scaling, with minimum value imputation for missing values and proteins expressed in <20% of cell lines removed in proteomics modalities. The top 1000 connections in the multi-omics networks were selected for visualization.

#### SH-SY5Y in vitro cultures, neuronal differentiation

SH-SY5Y cells (gift from Michael Whitehead) were kept in culture and expanded in growth medium (DMEM/F12, 10% FBS, 1% Penicillin/Streptomycin). Growth medium was refreshed every 4 days. After reaching 80-90% confluency, SH-SY5Y cells were washed with warm 1X PBS, detached with Trypsin-EDTA 0.05%, and spun down at 300 × g for 5 min. Next, SH-SY5Y cells were counted and seeded at a density of 10.4 × 10^3^ cells/cm^2^ in differentiation media (Neurobasal medium, 2% B27 supplement, 1% GlutaMAX, and 10 μM of all-trans retinoic acid [STEMCELL Technologies]). Differentiation media was replaced every other day until day 9 post-seeding, when SH-SY5Y neurons were exposed for 72h to fresh conditioned NIM from the control and PMS iNSC lines.

#### Simvastatin activation

Simvastatin requires activation through the opening of the lactone ring before it can be used in cell culture. To achieve this, we followed the procedure outlined by Merck. In a summary, we dissolved 8 mg of simvastatin (0.019 mM) in 0.2 mL of 100% ethanol. Subsequently, we added 0.3 mL of 0.1 N NaOH to the solution. The mixture was then heated at 50°C for 2 hours and later adjusted to a pH of 7.2 with HCl. The resulting solution was adjusted to a final volume of 1 mL using distilled water, aliquoted and stored at −80 °C until used to supplement cell culture media.

#### Generation of iNSC conditioned media for treatments

Control and PMS iNSCs were seeded in NIM with Y-27632 (10 µM) at a density of 100,000 cells/cm^2^ onto GFR-matrigel coated 6 well plates. Medium was refreshed the next day with NIM supplemented with 10 µM of simvatstatin or vehicle (ethanol) and cells were incubated for 48 hours. Medium was removed, ells were washed twice with 500 µL DPBS to remove simvastatin, and 1.5 mL of NIM was added to each well. After 24 hours conditioned medium (CM) was collected, spun at 300 × g for 5 minutes to remove debris and used immediately to treat SH-SY5Y neurons. For the concomitant treatment of SH-SY5Y neurons with ROCK inhibitor, 50 µM Y-27632 was added to the CM from iNSCs before it was placed onto the neurons. After 72 hours of CM exposure SH-SY5Y cells were collected for apoptosis and neuronal morphology assays.

#### Apoptosis assay

To evaluate neuronal death SH-SY5Y cells were seeded and differentiated onto 13 mm diameter coverslips. Three coverslips were seeded per condition. After CM treatment cells were fixed with 4% paraformaldehyde, washed twice with PBS for 5 minutes, and blocked for 1 hour at room temperature in 0.1% PBST with 10% normal goat serum. Coverslips were then incubated at 4°C overnight with cleaved caspase 3 antibody (Cell Signaling,1:1000) in blocking buffer. Next the coverslips were washed three times with 0.1% PBST for 5 minutes, incubated with the appropriate secondary antibody (1:1000) in blocking buffer for 1 hour at room temperature, and then washed with PBS three times for 5 minutes. Lastly, the coverslips were counterstained with DAPI then mounted. 63x images pictures of 3 ROIs per coverslip were taken on a Leica DMI400B microscope. Images were batched analyzed using Fiji software; the pipeline included fluorescent signal identification by minimum error thresholding followed by the measurement of the mean gray value. The values from 3 ROIs were averaged out for each coverslip, and the resulting values were again averaged out to generate the mean fluorescence intensity (MFI) of each condition (e.g., neurons exposed to CM from PMS iNSC line A, neurons exposed to CM from control iNSC line B).

#### Assessment of neurite length

SH-SY5Y cells were seeded and differentiated onto 6 well plates. Three wells were seeded per condition. After CM treatment SH-SY5Y neurons were imaged using an Echo Rebel microscope (Discover Echo) at 10x. Phase contrast images of 5 ROIs per biological replicate from each condition were taken. Images were converted to 8-bit and contrast was adjusted to make neurites easily visible using FIJI software. Data was quantified by a semi-automatically tracking NeuronJ plugin^81^ from an average of >15 neurites per ROI. The values from 5 ROIs were averaged for each well, and the resulting values were averaged to generate the mean neurite lengths of each condition.

#### Quantification and Statistical Analysis

ImageJ was used to analyze immunofluorescence and immunoblot images, and the detailed procedure is described in the ICC and immunoblot sections respectively. GraphPad Prism 10 for macOS was used to calculate statistics for non-omics data. Data was analyzed with the statistical test indicated in each figure. In the figure legends levels of significance (*p<0.05, **p<0.01, ***p<0.001, ****p<0.0001), along with statistical test and n are included. Metabolomics and lipidomics datasets were pre-processed using per-total normalization and feature standardization, on mean and standard deviation. Principal component analysis (PCA) and PLS-DA analysis were performed using prcomp() (stats v4.2.1 R package) and plsda() (mixOmics v6.22.0 R package (https://doi.org/10.1371/journal.pcbi.1005752)) functions, respectively. Disease and treatment groups were compared using Welch’s t-test with Benjamini-Hochberg multiple testing correction. Proteomics datasets were pre-processed using minimum value imputation of missing values, followed by median normalization. Further exploratory analyses were performed using bulkAnalyseR v1.1.0 (https://academic.oup.com/bib/article/24/1/bbac591/6965538) with differential expression calls using edgeR v3.40.2 (https://doi.org/10.1093/bioinformatics/btp616) with Benjamini-Hochberg multiple testing correction, and enrichment analysis performed on differentially expressed proteins using gprofiler2 v0.2.2 (https://doi.org/10.1093/nar/gkz369); the set of all genes expressed in matching RNAseq data was used as the background, only proteins expressed in more than 20% of cell lines were considered and FDR multiple testing correction was also performed.

