## Supplemental Figures for "Increased Cholesterol Synthesis Drives Neurotoxicity in Patient Stem Cell-Derived Model of Multiple Sclerosis"

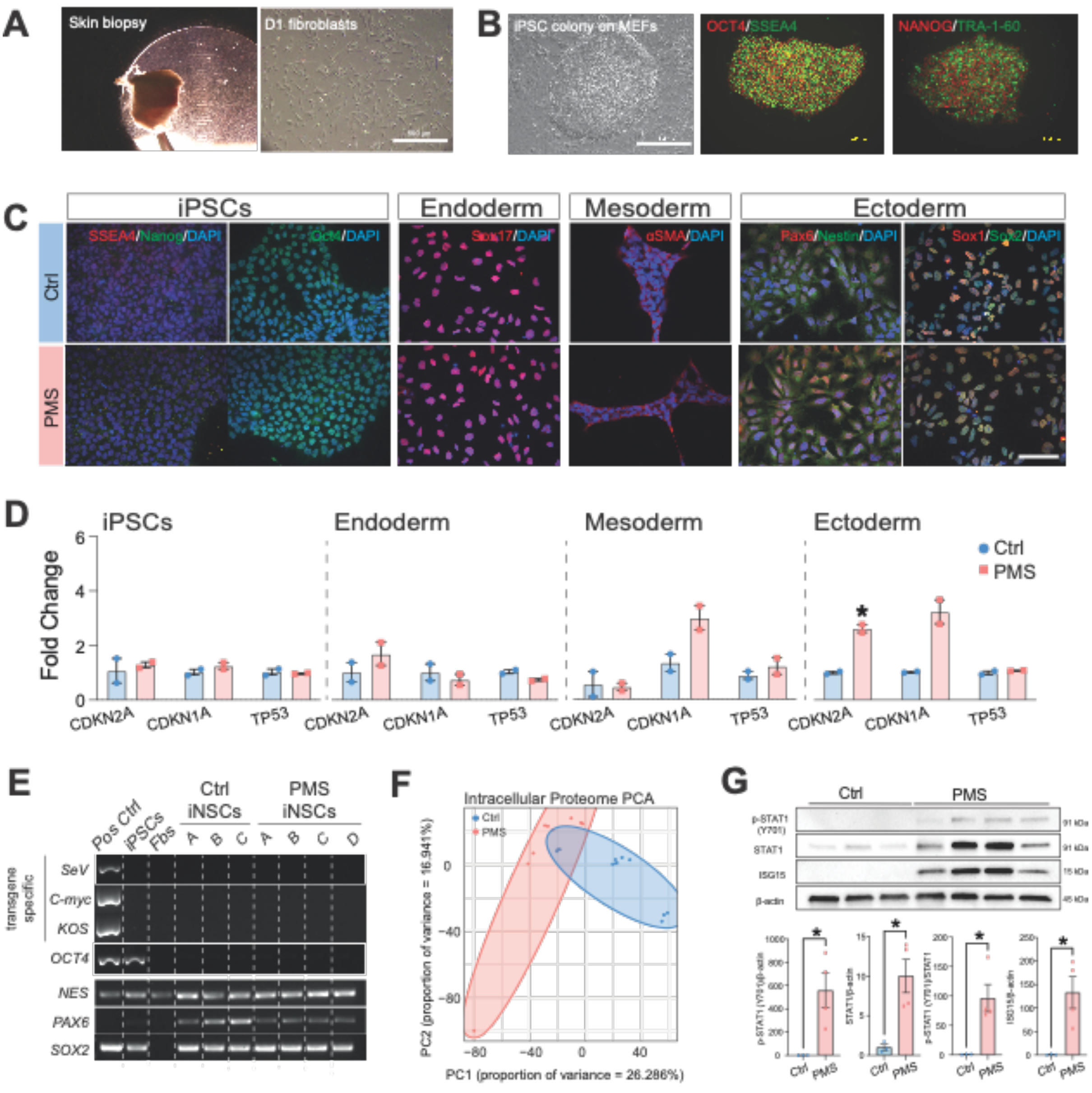


Ionescu, Nicaise et al. Figure S1

### Figure S1. Expression of senescence-associated genes and inflammatory proteins

(**A**) Phase contrast images of skin punch biopsy and resulting fibroblasts after one day in culture.

(**B**) Phase contrast and immunocytochemistry of iPSC markers.

(**C**) Immunocytochemistry for accepted iPSC and trilineage markers (endo-, meso-, ectoderm). Scale bar: 100 μm.

(**D**) mRNA expression of senescence-associated genes.

(**E**) RT-PCR gel of transgenes from Sendai virus reprogramming, pluripotency genes, and accepted NSC genes. Fbs, fibroblasts.

(**F**) PCA of the intracellular proteome.

(**G**) Representative western blot and quantification for p-STAT1 (Y701), STAT1, and ISG15.

Data in **D** and **G** are mean values ± SEM. Experiments were done on n= 2-3 Ctrl and n= 2-4 PMS iNSC lines, each performed in n ≥ 2 replicates. *p≤ 0.05, unpaired t-test.

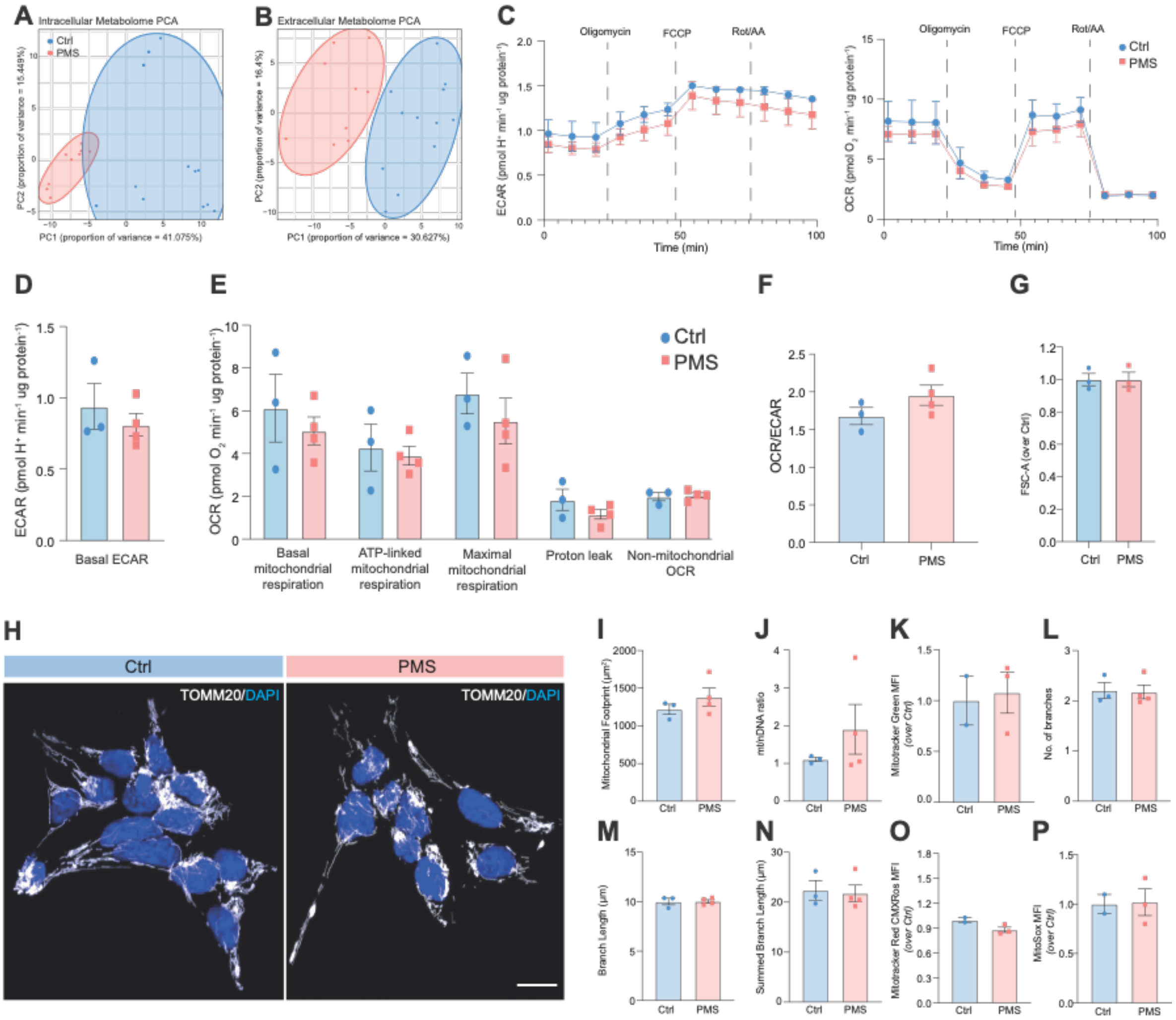


Ionescu, Nicaise et al. Figure S2

**Figure S2. Metabolic features of iNSC and fibroblast lines from people with PMS and Ctrl counterparts**

(**A** and **B**) PCAs of the intracellular and extracellular metabolome of iNSCs.

(**C**) ECAR and OCR over time during a mitochondrial stress test in the presence of glucose, pyruvate, and glutamine in fibroblasts.

(**D**) Basal ECAR as in **C**.

(**E**) Quantification of basal mitochondrial respiration, ATP-linked mitochondrial respiration, spare mitochondrial respiration, mitochondrial respiration linked to proton leak, and non-mitochondrial OCR using the mitochondrial stress test as in **C**.

(**F**) OCR/ECAR ratio of iNSCs.

(**G**) Flow cytometry-based quantification of cell size in iNSCs using forward scattering profiles.

(**H**) Immunocytochemistry for TOMM20 in iNSCs. Scale bar: 25 μm.

(**I**) Quantification of mitochondrial footprint using TOMM20 immunocytochemistry and the MiNa ImageJ plugin as in **H**.

(**J**) Quantification of mitochondrial copy number in iNSCs by qPCR. The number of copies of mitochondrial gene MT-ND2 is normalized to the number of copies of nuclear gene RNaseP.

(**K**) Flow cytometry-based quantification of mitochondrial mass in iNSCs using MitotrackerGreenMFI.
(**L**) Quantification of mitochondrial network branches as in **H**.
(**M** and **N**) Quantification of mitochondrial network branch length and summed branch length as in **H**.
(**O**) Flow cytometry-based quantification of mitochondrial membrane potential in iNSCs using MitotrackerRed CMXRos and normalized to mitochondrial mass.
(**P**) Flow cytometry-based quantification of mitochondrial superoxide levels in iNSCs using MitoSoX.
Experiments were done on n= 2-3 Ctrl and n= 2-4 PMS iNSC lines (**A**, **B**, **F, G** and **H**-**P**), or same number of fibroblast cell lines from Ctrl and PMS lines (**C-E**), each performed in n ≥ 3 replicates. Data in **C**-**G and I**-**P** are mean values ± SEM. p≥ 0.05, unpaired t-test.

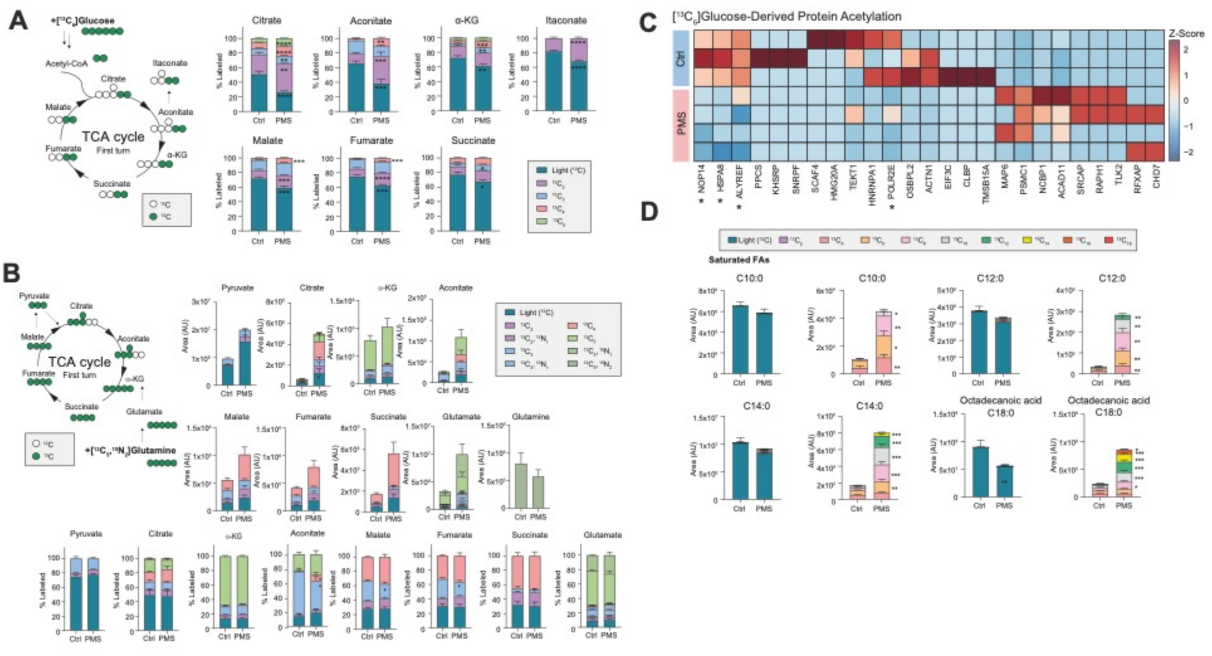


Ionescu, Nicaise et al. Figure S3

### Figure S3. [$^{13}C_6$]glucose or [$^{13}C_5$,$^{15}N_2$]glutamine tracing in PMS and Ctrl iNSCs

(**A**) Intracellular enrichment in TCA cycle at 24 h following [$^{13}C_6$]glucose tracing. Data is presented as proportional enrichment of total metabolite pools.

(**B**) Intracellular enrichment in TCA cycle at 24 h following [$^{13}C_5$,$^{15}N_2$]glutamine tracing. Data is presented as peak areas (arbitrary units) and as proportional enrichment of total metabolite pools.

(**C**) Protein acetylation following 6 hours of [$^{13}C_6$]glucose tracing.

(**D**) Intracellular enrichment in saturated fatty acids at 24 h following [$^{13}C_6$]glucose-tracing. Data is presented as peak areas (arbitrary units).

Experiments were done on n=3 Ctrl and n= 4 PMS iNSC lines, each performed in $n \geq 3$ replicates. Data in **A, B,** and **D** are mean values ± SEM. *$p \leq 0.05$, **$p \leq 0.01$, ***$p \leq 0.001$, ****$p \leq 0.001$, unpaired t-test.

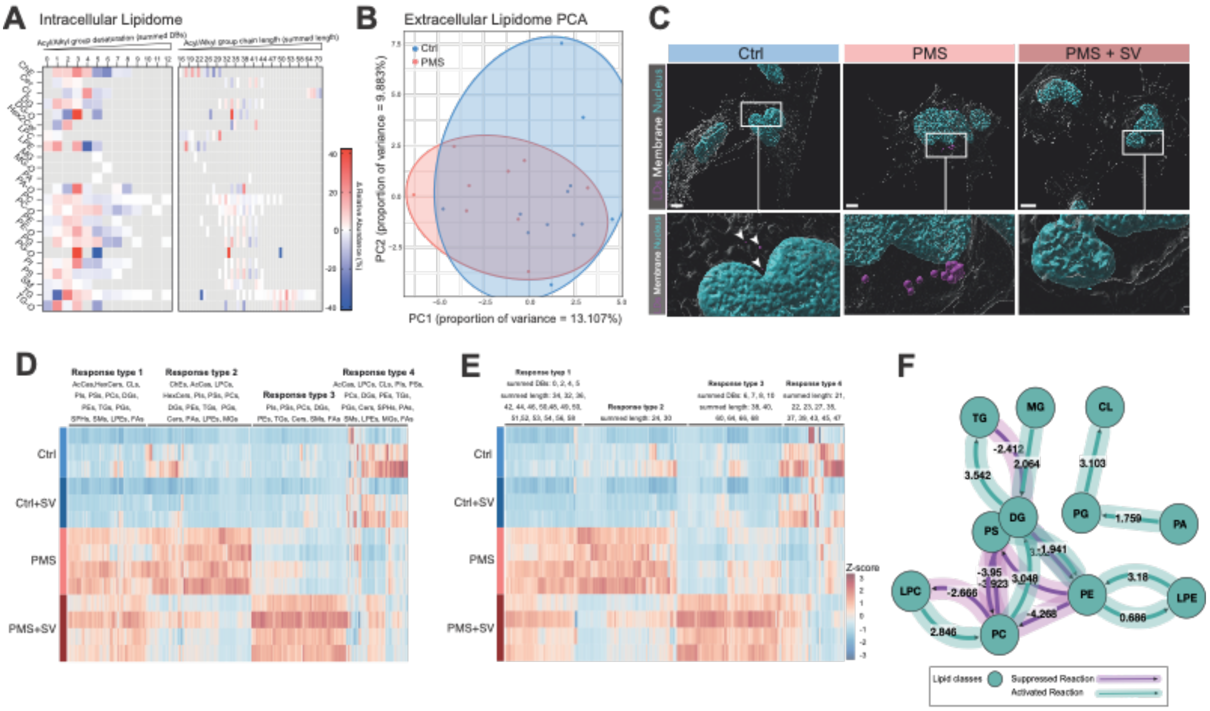


Ionescu, Nicaise et al. Figure S4

### Figure S4. Lipidomic profiling

(**A**) Structural changes in lipid species in PMS iNSC (*vs.* Ctrl iNSCs). Relative abundance is expressed as percent change in acyl/alkyl group desaturation (as summed double bonds [DBs]) and in acyl/alkyl group chain length (as summed length).

(**B**) PCA of the extracellular lipidome.

(**C**) 3D reconstructed high resolution images of LipidSpot 610[+] LDs and BioTracker 555-stained cytoplasmic membranes. Scale bars: 5 µm. Arrowheads indicate three small LDs in Ctrl iNSCs.

(**D** and **E**) Hierarchical clustering of intracellular lipid species (**D**) and deconvoluted lipid species substructures (**E**). Clustering in **D** shows the distribution of lipid classes across the four identified responses.

(**F**) BioPAN analysis of the top activated and suppressed intracellular lipid class reactions induced by inhibition of HMG-CoA reductase with SV in PMS iNSCs. A reaction is considered significantly modified at a level of $p \leq 0.05$ (corresponding to $Z \leq 1.645$).

Experiments were done on n= 3 Ctrl and n= 4 PMS iNSC lines, each performed in $n \geq 3$ replicates.

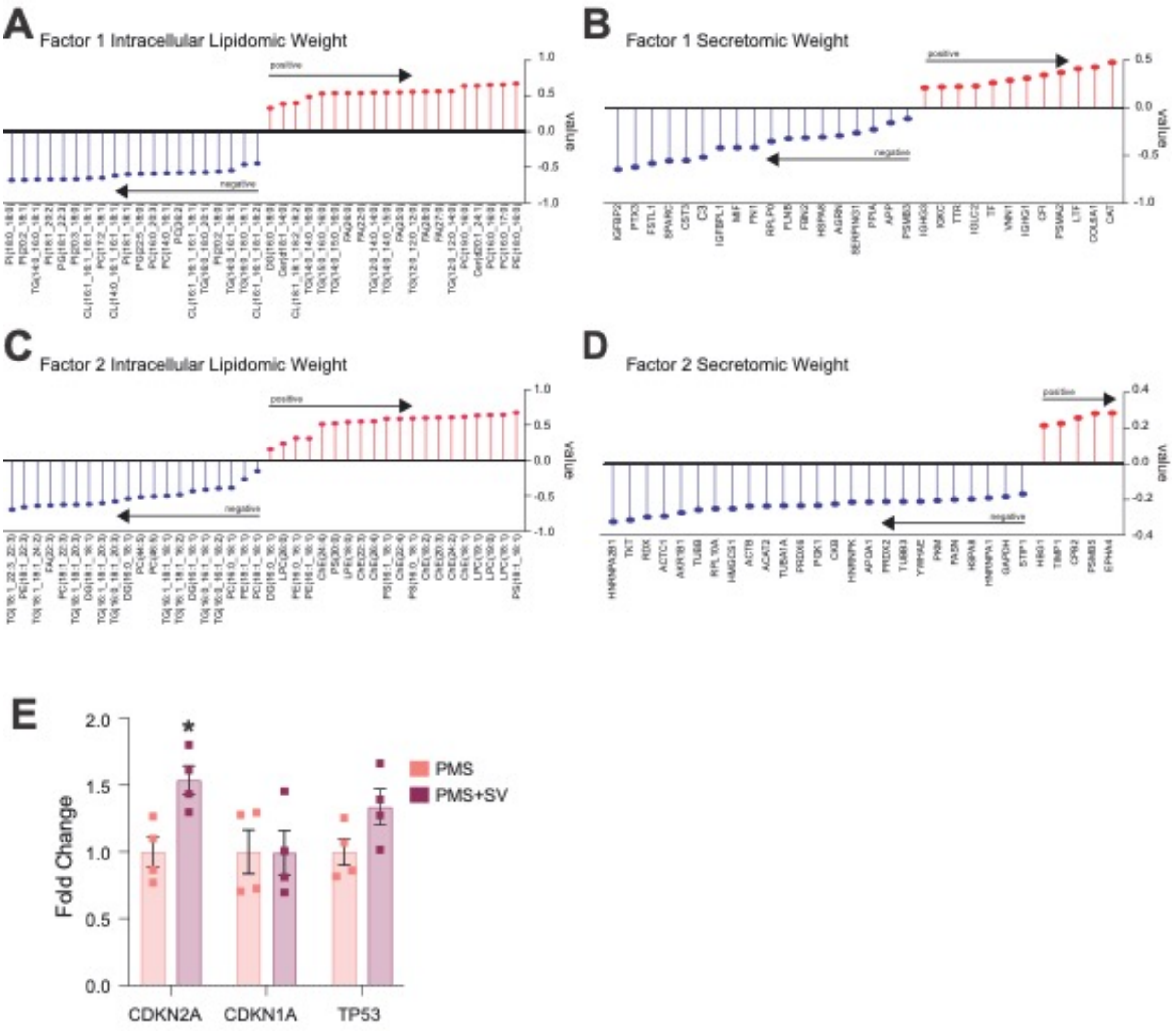


Ionescu, Nicaise et al. Figure S5

**Figure S5. Co-regulated features revealed by MOFA**

(**A** and **B**) Top intracellular lipidomic and secretomic features contributing to latent Factor 1.

(**C** and **D**) Top intracellular lipidomic and secretomic features contributing to latent Factor 2.

(**E**) mRNA expression of senescence-associated genes after SV treatment in PMS iNSCs.

Data in **E** are mean values ± SEM. Experiments were done on n= 3 Ctrl and n= 4 PMS iNSC lines, each performed in n ≥ 3 replicates. *p≤ 0.05, unpaired t-test.
